# The menstrual cycle through the lens of a wearable device: insights into physiology, sleep, and cycle variability

**DOI:** 10.1101/2025.09.11.675620

**Authors:** Alexander Gonzalez, Johanna J. O’Day, Sarah C. Johnson, Jeongeun Kim, Summer Jasinski, Kristen Holmes, Scott L. Delp, Jennifer L. Hicks

## Abstract

Women on average have 450 menstrual cycles in a lifetime, but we lack a characterization of physiological biometrics across the cycle and lifespan. We analyzed 1.2 million days of data from 2,596 women who logged 42,759 menstrual cycles and wore a device that collected sleep and biometric data including resting heart rate (RHR), heart rate variability, respiratory rate, skin temperature, and blood oxygen saturation level. We generated novel quantifications of daily biometrics across ages and cycle lengths, finding that cycle length is strongly associated with how much cardiorespiratory metrics vary across the cycle. We observed greater cycle variability for participants who slept 6 versus 8 hours. A within-participant natural experiment showed that decreased sleep resulted in biometric changes regardless of cycle phase (e.g., RHR increased 1.2% with a 10% decrease in weekly sleep duration). These results lay a foundation to better understand and optimize female health and performance.

## Introduction

Many women report changes in physiology, sleep, and athletic performance over the menstrual cycle^1–3^. Yet there is a marked lack of understanding of how physiological biometrics change across the cycle, how these patterns change with age, and how variations in physiology across the cycle relate to behavior and performance.

The *length* of a menstrual cycle and the *variability* of cycle length are potential biomarkers for female health^4^. Individuals who reported consistently high cycle variability were more likely to report negative menstrual symptoms^5^, while cycle irregularity (missing 3 or more consecutive menstrual cycles per year) has been used in a clinical scoring of Relative Energy Deficiency in Sport^6^. Long and irregular menstrual cycles have also been associated with other serious long-term health outcomes, including cancer, diabetes, cardiovascular disease, fracture incidence, and premature mortality^4^. However, it remains unclear how behaviors, such as reduced sleep, affect cycle length and cycle variability.

Consistent cyclic variation in basal body temperature^7^ and skin temperature^8^ has been shown to relate to changes in reproductive hormones and thus is used by women to track fertility status^7,9^. Researchers have also found relationships between biometrics, such as resting heart rate and phases of the menstrual cycle^10–16^, pregnancy^17,18^, and age^16^. Digital logging through smartphone applications and wrist-worn and finger-worn devices makes it more convenient to track menstruation, physiological biometrics, and behaviors like sleep and workout activity.

Studies using digital health tools have replicated findings from previous in-lab studies^19–22^, showing that follicular phase length, and resulting cycle length, decrease with age^23–25^. Digital health studies have also confirmed previous observations that skin temperature^8,12,26^ and resting heart rate^12,14,16,26^ are lowest in the follicular phase and rise in the luteal phase. Even with these recent advances, biometrics at a daily resolution have not been reported across ages and cycle lengths, nor have their independent impacts been assessed.

Alongside the biometric and hormonal fluctuations across the cycle, before or during the menstrual week, women report lower motivation to train^3^, higher rates of “sadness”^27^, and lower sleep quality^28,29^. However, prior studies aiming to identify cycle-related differences using objective measures of sleep^29^ and performance^30^ have been inconclusive. These reviews highlight common limitations of past studies—namely the small number of participants and reliance on few menstrual cycles per participant—restricting their ability to capture trends amid high individual variability. Thus, it remains unclear how reduced sleep affects menstrual cycle characteristics, such as cycle length and physiological biometrics, in real-world settings.

Equipped with a large-scale, longitudinal, wearable-based dataset, we addressed several gaps in menstrual cycle research. First, we identified how sleep duration impacts menstrual cycle variability, leveraging participant-reported menstrual cycles together with objective and passively recorded sleep behaviors. Second, we established a detailed normative profile of physiological biometrics, including resting heart rate, heart rate variability, skin temperature, respiratory rate, and blood oxygen saturation, throughout the menstrual cycle. We further modeled the independent effects of age and cycle length on these biometric profiles, offering quantifications previously absent in the literature. Finally, we analyzed how sleep changed across the menstrual cycle and quantified the impact of changes in sleep on resting heart rate and the other biometrics.

## Results

### Dataset description and verification of known age-cycle relationships

We analyzed 1,298,555 days and 42,759 menstrual cycles from 2,596 participants aged 18-50 years old who had not reported using hormonal contraception or being pregnant or in peri/menopause during the analysis period (age: 33.7 mean ± 6.5 S.D., body mass index (BMI): 24.8 ± 4.4, Supplementary Table 1; see Methods for additional details about participant inclusion/exclusion). Participants regularly wore a device, typically on the wrist, that recorded daily metrics including sleep onset and duration (sleep duration: 7.2 ± 0.6 hours/night). Additionally, the device recorded nightly measures of resting heart rate (RHR), heart rate variability (HRV), respiratory rate (RR), skin temperature (Temp.), and blood oxygen saturation level (Blood O2). Participants used a connected smartphone application to log their menstruation status. These logs were then used to compute cycle lengths.

We reproduced known relationships between cycle length, cycle variability, and age, giving confidence in the validity of the self-reported menstrual cycles. The average cycle length was 28.4 ± 2.4 days, in agreement with prior reports and clinical standards^31^. We observed a decrease in cycle length with increasing age (Figure 1a), in agreement with clinical studies^19–21^ and large-scale observational digital health studies^23–25^. Cycle length decreased from 29.1 days at age 24 years to 26.9 days at age 44 (difference 2.3 [95% CI: 2.0 - 2.5]) when controlling for BMI (Figure 1a, see Figure S1 for relationship between cycle length and BMI; see Methods for statistical model details).

**Figure 1.**
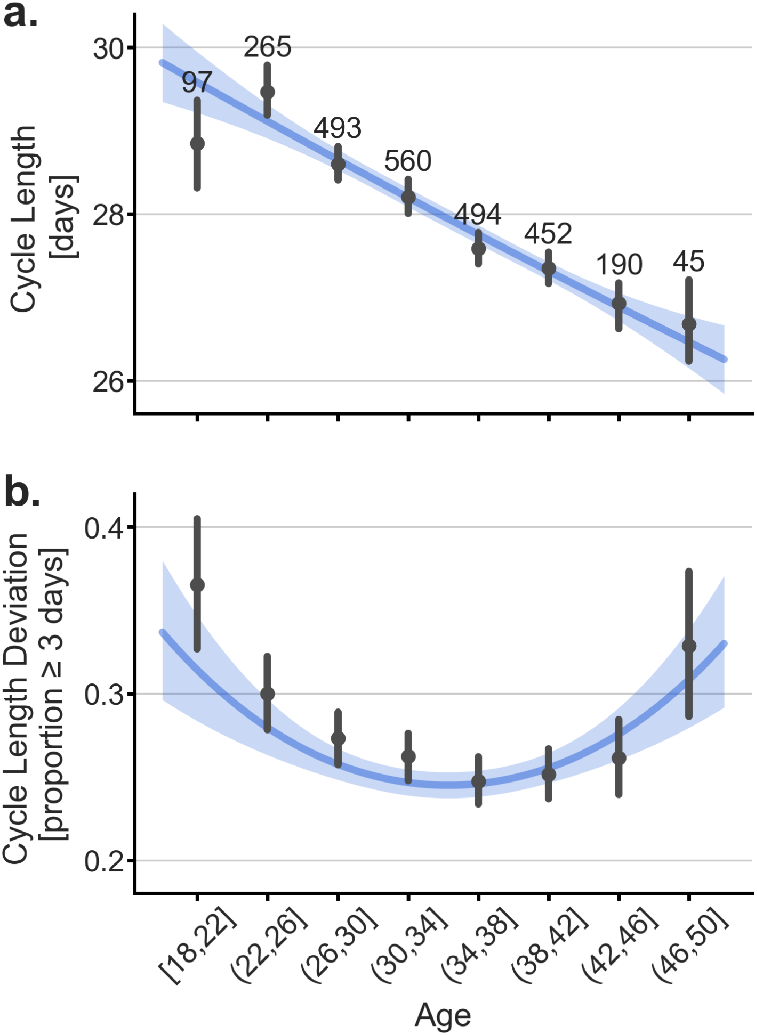
Menstrual cycle length and menstrual cycle length deviation vary with age. **(a)** Cycle length decreases with age. Each point represents the mean cycle length across participants for a given 4-year age bin from 18 - 50 years (x-axis). Annotations above the points indicate the number of participants per age bin and error bars indicate the bootstrapped 95% CI of the mean. The blue line represents the conditional expectation of cycle length from a Generalized Estimating Equation (GEE) model relating cycle length to age, with the shaded region representing the 95% CI. **(b)** Cycle length deviation (proportion of cycles deviating from a participant’s median cycle length by 3 days or more) versus age followed a U-shaped pattern, as confirmed with a GEE model (blue line). Points and error bars as in (a) with cycle length deviation instead of cycle length.

There was a U-shaped relationship between cycle length deviation and age (Figure 1b; see Methods for statistical model details). Across the cohort, 27% of cycles deviated by three days or more from a participant’s median. The most consistent cycle lengths were at age 33 [95%CI: 31 - 35], where 1 in 4 cycles (0.25 [95%CI: 0.24 - 0.25]) deviated by three or more days from the median. At age 24, the frequency of cycle length deviation was slightly higher at 0.28 [95%CI: 0.26 - 0.30]. These relationships were also observed with other definitions of cycle variability, including the standard deviation of cycle length (Figure S2). These findings are in alignment with previous reports on cycle variability trends with age^19–21,23–25,32^.

### Sleep and cycle length

To explore the relationships between cycle characteristics and sleep patterns, we used Generalized Estimating Equations (GEE) models to relate cycle lengths to sleep duration and sleep duration variability. The models controlled for age, BMI, sleep onset, and workout patterns (Figure S3; see Methods for model details).

Sleep duration had a strong association with deviations in cycle length (Figure 2b). Participants were more likely to have a cycle length that deviated by three or more days from their median when sleeping 6 hours (cycle deviation proportion = 0.29 [95%CI: 0.28 - 0.31]) compared to 8 hours (0.25 [95%CI: 0.24 - 0.26]). That is, there was a 1.3 times greater odds ratio (OR) of a cycle length deviation for sleeping 6 versus 8 hours [95%CI: 1.2 - 1.4]. Sleep duration had only a weak association with mean cycle length (e.g., cycle lengths were 0.4 days shorter for those sleeping 6 versus 8 hours [95%CI: 0.1 - 0.6], Figure 2a).

**Figure 2.**
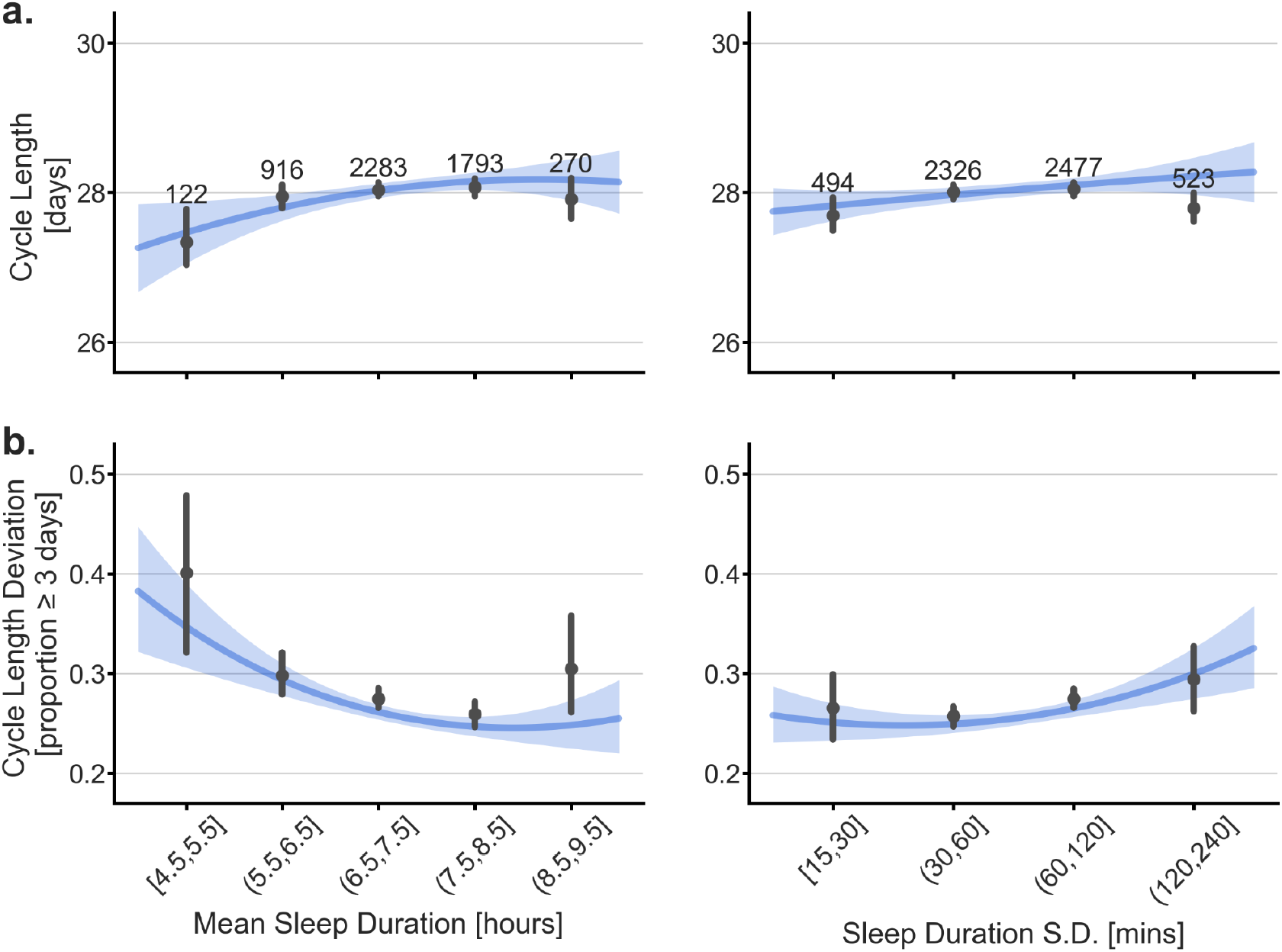
Population level relationships between sleep duration and cycle length. **(a)** Cycle length was weakly associated with mean sleep duration (left) and sleep duration variability (right). Left, for each 1-hour sleep duration bin (x-axis), the points represent the average cycle length across participants who had cycles with an average sleep duration in that bin. Right column same as the left but with sleep duration standard deviation (S.D.) bins (x-axis). Annotations above the points indicate the number of participants with at least one cycle in that bin, with error bars indicating the bootstrapped 95% CI of the mean. The blue line represents the conditional expectation from a GEE model fitted to the data with population means for covariates (e.g., age=33, BMI=25) and with the shaded region representing the conditional 95% CI. **(b)** Cycle length deviation had a strong, non-linear relationship with sleep duration and sleep duration variability. Relationships quantified through GEE models with fixed covariates as in (a) and cycle length set to 28. Points and error bars as in (a) except with cycle length deviation instead of cycle length. We observed statistically significant differences in cycle length deviation for mean sleep durations below 7.3 hrs (min at 8.4 hrs [95% CI: 7.3 - 9.5 hrs]) and sleep duration variabilities above 55 minutes (min cycle length variability at 32 minutes [95% CI: 19-55 minutes]).

In addition to our population-level findings, we examined a subset of participants who changed their sleep behavior during the measurement period (N=308). These participants had at least one cycle with an average sleep duration between 5.5 and 6.5 hours and one with an average between 7.5 and 8.5 hours. We found a similar odds ratio for higher cycle length deviation with shorter sleep duration to our population level analysis, although the relationship was not significant at the alpha=0.05 level (OR=1.2 [95%CI: 0.9 - 1.6]).

Variability in sleep duration has also been related to health outcomes^33^. Using a similar GEE population level modeling approach, we found that greater variability in sleep duration was associated with slightly longer cycles (see Methods for model details). For example, a cycle with a sleep duration variability of 120 minutes led to a 0.3 day increase in cycle length [95% CI: 0.1 - 0.5], relative to a cycle with 30 minutes of variability (Figure 2b). This increased variability in sleep duration also yielded 1.2 times greater odds of a cycle length deviation of three or more days [95%CI: 1.1 - 1.3].

A within-participant analysis confirmed this relationship between increased sleep variability and more variable cycle lengths (OR=1.3 [95%CI: 1.1 - 1.7]). This sub-analysis consisted of 813 participants who had at least one cycle with high sleep duration variability (85 - 170 minutes) and one cycle with low sleep duration variability (21 - 42 minutes).

### Biometrics across the menstrual cycle as a function of age and cycle length

We used the daily-resolution physiological biometrics to characterize fluctuations across the menstrual cycle. We first verified that the dataset could reproduce known age-related trends in biometrics (Figure S4). For example, we reproduced the prior findings that HRV decreases with increasing age^34^ (Figure S4) and increasing BMI^35^ (Figure S5). Together, these results gave us confidence in using the wearable-based dataset to study biometrics across the cycle.

All biometrics but HRV dipped during menstruation and were highest before the next menstrual cycle onset (Figure S6). HRV showed the opposite pattern, with a peak during menstruation.

The cardiovascular metrics (RHR, HRV, and RR) were significantly correlated with each other throughout the cycle (|r|>0.25, based on a Vector Autoregressive model that accounted for autocorrelation effects, see Methods for model details; Figure S7a). Moreover, RHR explained 49% of the future variability in HRV and 18% of the variability in RR, but only 3% of the variability in skin temperature (based on variance decomposition analysis; Figure S7b).

We generated normative temporal profiles of biometrics across ages (Figure 3a) and cycle lengths (Figure 3b) by fitting Generalized Additive Models (GAM) to the cohort data (see Methods for model details). The GAM temporal profiles allowed us to identify the population-level timings of minima, zero-crossing, and maxima for each biometric across the cycle (Figure S8). The timing of these fluctuations in biometrics varied by cycle length. For RR and skin temperature, the timing of the minima was strongly influenced by cycle length (e.g., RR: minima at day 13 for a 34-day-cycle compared to day 8 for a 24-day-cycle; fixed age at 32). In contrast, the RHR minima and HRV maxima were reached at day 6 for a 34-day-cycle vs day 4 for a 24-day-cycle. Aging shifted both the minima and maxima timings earlier by fewer than 2 days for RHR and HRV. Blood O2 showed some cyclicality (Figure 3), but the effects of age and cycle length were relatively small and highly variable (model explained deviance = 1%).

**Figure 3.**
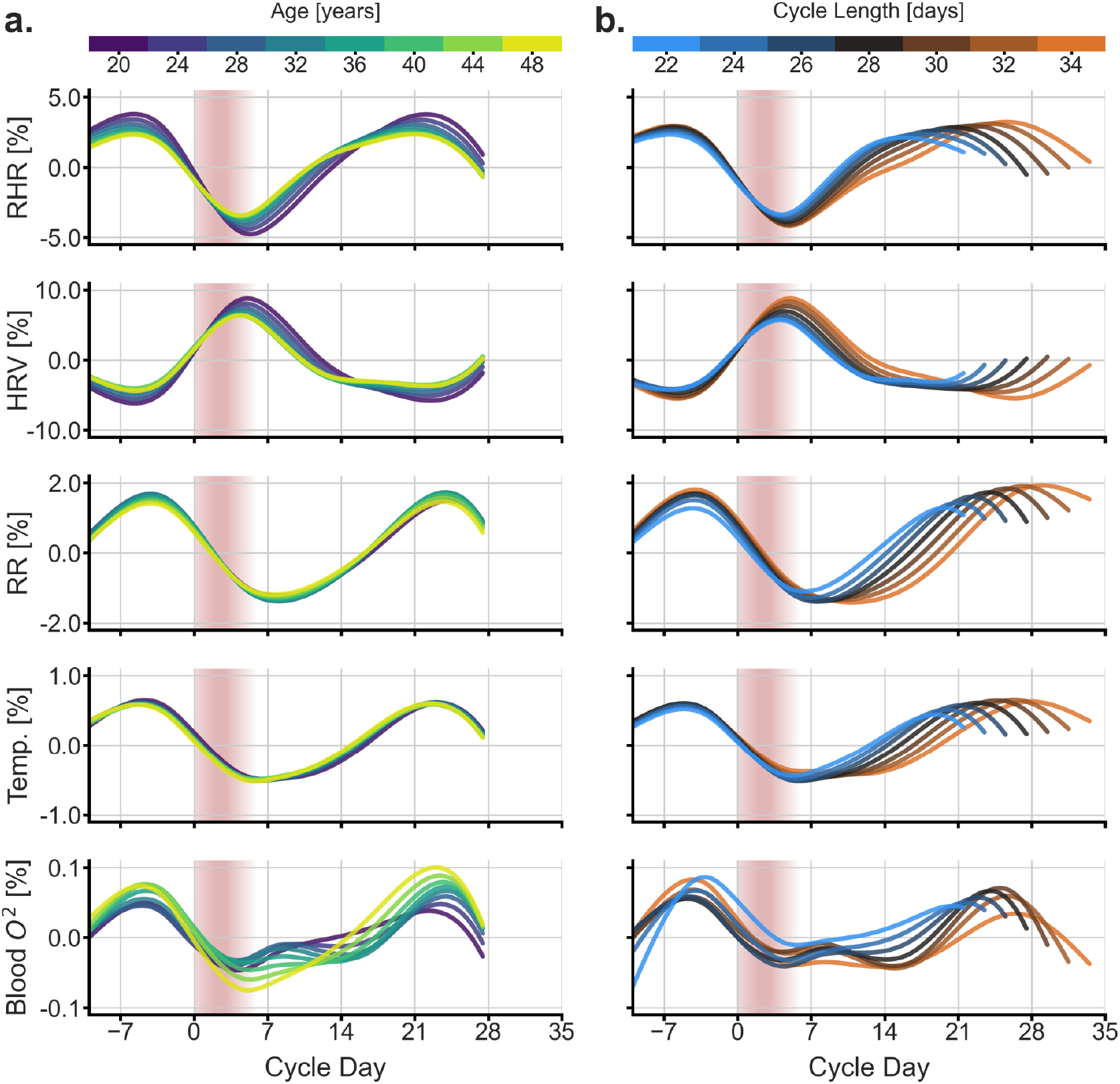
Biometrics as a function of age and cycle length. We generated temporal biometric profiles across a variety of **(a)** ages (with fixed cycle length of 28 days) and **(b)** cycle lengths (with fixed age of 32 years) using Generalized Additive Models fitted to each normalized biometric. Biometrics: resting heart rate (RHR; model deviance explained D^2^ = 31%), heart rate variability (HRV; D^2^ = 14%), respiratory rate (RR; D^2^ = 31%), skin temperature (Temp.; D^2^ = 23%), and blood oxygen saturation level (Blood O^2^; D^2^ = 1%). Biometric curves were generated for ages in 20-48 years in 4-year steps and cycle lengths 22-34 days in 2 days steps (legend). Typical menses is depicted in red, starting with menstrual cycle onset (day zero).

RHR and HRV showed an age-related decrease in the range (peak – trough) across the cycle as computed from their GAM profiles (Figure S9a). For example, the range in HRV across the cycle decreased from 13.3% of the mean HRV value to 10.0% over 20 years (age 24 vs 44 with a 28-day cycle; difference=3.3% [95%CI: 2.9 - 3.6%]). Conversely, the range of other biometrics changed minimally with age (e.g., RR range decreased from 2.9% to 2.8% over 20 years). As cycle length increased, the range of cardiorespiratory biometrics across the cycle increased. For example, the HRV range increased from 9.4% to 14.3% with a 10 day-cycle increase (cycle length 24 vs 34 at age 30; difference=4.9% [95%CI: 4.4 - 5.4%]), and the RR range increased from 2.7% to 3.3% (cycle length 24 vs 34 at age 30; difference=0.7% [95%CI: 0.6 - 0.8%]). The range of skin temperature and blood oxygen saturation level did not significantly differ by age or cycle length.

The profiles from the GAM reflect the population-level waveforms across ages and cycle lengths. At the individual-level, within-cycle biometric ranges were expectedly greater than the population-level normative profiles. For an individual cycle, HRV varied on average by 38% of a participant’s mean value, RHR by 14%, RR by 6%, Temp. by 3%, and Blood O2 by 2% (Supplementary Table 2 and Figure S9 for a breakdown by age and cycle length).

### Sleep and physiology across the cycle

Given anecdotal reports of sleep varying across the menstrual cycle^36^, we quantified the percentage change in sleep duration during a given cycle week (premenstrual, menstrual postmenstrual). We focused on time periods following at least three weeks of stable sleep (7-8 hours on average) and generated distributions of changes in sleep duration by menstrual cycle week (Figure 4a). Using a similar GEE modeling approach as above (see Methods for additional details), we found that decreases of 10% or more in sleep duration were more prevalent in the premenstrual week (premenstrual to menstrual OR: 1.39 [95%CI: 1.14 - 1.68], premenstrual to postmenstrual OR: 1.34 [95%CI: 1.11 - 1.62], menstrual to postmenstrual OR: 1.03 [95%CI: 0.85 - 1.27]).

**Figure 4.**
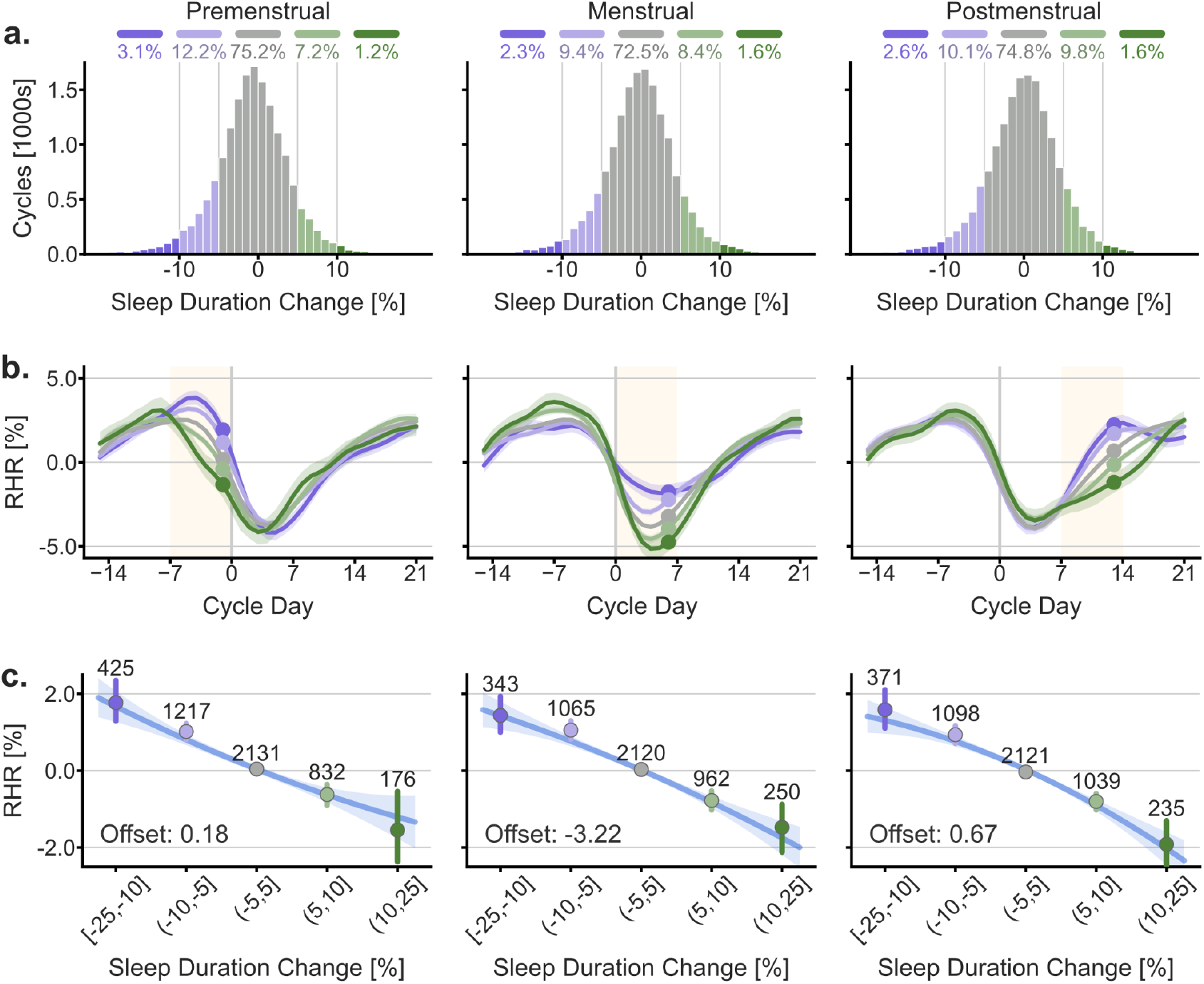
Changes in sleep duration relate to changes in resting heart rate consistently across the menstrual cycle. **(a)** The distribution of sleep duration changes as a function of cycle week (left - premenstrual week: # cycles = 16,441, mean=-1.1%; center - menstrual week: 17,221, m=- 0.4%; right - postmenstrual week: 16,518, m=-0.3%). Changes are relative to time periods with 7-8 hours of average sleep duration in the prior three weeks. Colored regions indicate percentage threshold of sleep duration changes (e.g., −10%, light purple representing ∼42-48 minutes of decreased sleep for that week). Grey “no change” regions highlight cycles where participants did not change their sleep duration by more than 5% during that week compared to the preceding three weeks. Percentage annotations represent the fraction of cycles that had a given level of sleep duration change for that week. **(b)** Resting heart rate (RHR) by menstrual cycle day for varying amounts of sleep duration change. Colored lines represent the average resting heart rate from cycles corresponding to the sleep duration change levels from panel (a). Resting heart rate lines include shaded 95% CI. The shaded orange region on each panel represents the week of the cycle where the sleep duration change took place; there were no additional constraints on sleep duration past that week. **(c)** RHR linearly scales with changes in sleep duration across the menstrual cycle. Points correspond to the average RHR on the last day of the cycle week when the sleep change occurred (points in the curves of panel (b); day −1 for premenstrual, day 6 for menstrual, day 13 for postmenstrual). Annotations above each point represent the number of participants with at least one cycle at that sleep duration change and error bars indicate the bootstrapped 95% CI of the mean. The blue line represents the predicted RHR at different sleep duration change levels, from a Generalized Estimating Equation model fit to the data, using the population mean of the covariates (age, BMI, sleep duration). We shifted the RHR y-axis for each menstrual cycle week to reflect the change from the no-change in sleep duration condition (|change|<5%; offset annotation).

Next, we sought to identify if changes in sleep duration were associated with changes in biometrics and if these associations were different across the menstrual cycle (Figure 4b). A GEE model revealed an inverse, linear relationship between RHR and changes in sleep, such that a 10% decrease in sleep duration resulted in a 1.2% increase [95%CI: 0.9 - 1.4] in RHR (Figure 4c; see Methods for model details). We found no significant differences between cycle weeks in how RHR responded to changes in sleep duration (interaction χ_4_=4.5, p=0.3). We observed similar relationships between sleep duration and biometric patterns for HRV, RR, and Blood O2 (Figure S11). Skin temperature tended to increase with decreasing sleep duration, but the pattern varied depending on the menstrual cycle week (Figure S11d, χ_4_=10.5, p=0.03). In the premenstrual week when skin temperature was high, decreases in sleep duration did not lead to further temperature increases. In contrast, during the menstrual week when skin temperature was low, increases in sleep duration did not lead to further temperature decreases.

## Discussion

Our analysis of a large-scale (42,759 cycles, 1.2 million days), real-world dataset collected from a wearable device and smartphone application revealed insights into physiology and sleep across the menstrual cycle. We found that shorter sleep durations were associated with increased variability in the length of the cycle. Analyzing the longitudinal dataset also revealed that nights with truncated sleep were more common in the premenstrual week. We discovered that decreases in sleep duration led to elevated resting heart rate, respiratory rate, skin temperature, and blood oxygen saturation, and decreases in heart rate variability. These relationships between sleep duration and biometrics were largely similar across the phases of the menstrual cycle. We share daily normative profiles of biometrics across the cycle which can serve as baselines for future studies, including novel quantifications of respiratory rate and blood oxygen saturation levels. Our findings align with and expand on existing literature, demonstrating the unique potential of longitudinal wearable-based data for female health research.

### Menstrual Cycle Variability and Sleep

By analyzing cycle variability across a continuum of sleep durations in the real world, our results identify a range of sleep durations (namely, those less than 7.3 hours) where risk for cycle variability is increased. This result goes beyond previous lab-based studies and qualitative reports that associate cycle variability with self-reported sleep difficulties^37^ and increased odds of shorter sleep (< 6 hours)^38,39^. We also found that when variability in sleep duration exceeded 55 minutes, there was an increased odds of observing more variable cycle lengths, consistent with other observations that sleep variability is related to adverse menstrual cycle symptoms^40,41^. Since poor sleep is also related to hormonal dysfunction^42^, it is unclear whether sleep behaviors are driving changes in hormonal fluctuations that then lead to greater cycle variability, or whether irregular hormonal fluctuations are driving decreases in sleep duration.

Thus, while we controlled for factors such as age, BMI, and activity level that can impact sleep and/or menstrual cycle characteristics, we are reporting associative, not causal relationships between cycle variability and sleep.

### Normative Physiological Profiles Across the Cycle

Despite the plethora of daily biometric information provided by wearable sensors, there is no detailed understanding of how these measurements are expected to change across the menstrual cycle. After separating the effects of age and cycle length on biometrics, we found an age-associated shrinkage in the range of RHR and HRV through the cycle, similar to a previous report^16^. We further found that longer cycle lengths were associated with larger fluctuations in RHR, HRV, and RR across the cycle. Breaking out the effect of cycle length goes beyond prior studies that looked at fewer biometrics and fewer time points across the cycle^7,8,10,12,14,15^. The normative biometric profiles across ages and cycle lengths could serve as a resource for women who closely monitor their biometrics and for researchers to know when and how to account for biometric fluctuations.

Cycle-length temporally modulated the waveform of all biometrics, a clear indication of cyclicality. This cyclicality of biometrics during the menstrual cycle can likely be attributed to characteristic hormonal fluctuations including the elevated estrogen in the follicular phase followed by elevated progesterone in the luteal phase. The relationship between the hormonal patterns and basal body temperature has been extensively quantified and used, with mixed results, to predict ovulation^43–45^. Skin temperature has similarly demonstrated an association with hormonal fluctuations^8,46^. We observed that the timing of skin temperature and respiratory rate returning to their mean values was similar to hormone-kit-derived follicular phase timings across cycle lengths from previous studies^23^. While the ovulation-related rise in progesterone and corresponding rise in temperature are well known^7^, our results highlight respiratory rate as an additional potential marker of hormonal transitions through the cycle.

We observed a cyclic pattern in cardiac metrics where RHR was at a minimum around menstruation and peaked before the next cycle onset, with HRV having the opposite pattern, as in Jasinski et al., 2024^16^. The change in cardiac metrics early in the cycle could potentially arise from the increased estrogen levels that drive vagal tone activity (increases HRV) and suppress the baroreflex response (decreases heart rate)^11,47^, but more research is needed. The multi-variate temporal analysis did not find a strong residual correlation nor future predictive power between respiratory rate and skin temperature. This suggests that the observed temporal associations between these two biometrics do not follow a one-to-one mapping with expected hormone levels, with additional research needed to untangle those relationships.

### Sleep Changes Across the Cycle and Related Physiological Impact

To date, it has been unclear if objective measures of sleep duration would align with self-reports of reduced sleep in the premenstrual phase^29^. The longitudinal measures in our data enabled us to create distributions of sleep duration changes by cycle phase. While 10% or more decreases in sleep duration were relatively rare during the premenstrual week (503 of 16,441 qualifying cycles in 425 participants), the odds were higher than in other phases of the cycle (at least 1.3x), in alignment with self-reports^28^. Notably, there were 176 participants in our study who had at least one instance of increased sleep during the premenstrual week (194 cycles), underscoring the fact that sleep through the cycle is idiosyncratic, as previously described through self-reports of sleep difficulty^36^. An important note is that we cannot ascertain whether sleep changes were driven by a conscious participant choice, changing underlying hormones (e.g., progesterone-related body temperature increases^2,48^) or other causes.

While the driving causes of changes in sleep in our population were not possible to ascertain, we observed that these sleep changes significantly affected all biometrics. We found that a 10% reduction in sleep duration (e.g., 42 minutes less sleep per night over a week) led to an average increase of 1.2% in RHR (e.g., ∼0.7 bpm for a RHR of 60 bpm, in alignment with prior sleep deprivation research^49,50^. Interestingly, we observed that the impact of sleep changes on cardiac biometrics did not vary according to the menstrual cycle phase. In contrast, there were differences across the cycle in the association between skin temperature and changes in sleep. For example, skin temperature showed a ceiling effect with decreased sleep during premenstrual week, potentially due to tight thermoregulation in women^51^. The practice of aligning behaviors with menstrual cycle phases to optimize health and performance, termed “cycle-syncing”, has grown in popularity, but lacks strong empirical support^52,53^. Our findings begin to provide insight on how changes in behaviors, like sleep, may (or may not) influence physiology differently across the cycle.

## Limitations

It is important to consider the characteristics of the cohort included in this study. For example, participants were likely more “active” and “health-conscious” than the average individual, as evidenced by their workout levels even at older ages (Figure S3) and consistent engagement with the wearable device and smartphone application. The high workout levels could also explain the lower average RHR in this cohort at ages above 40 (Figure S4) compared to other large population studies^34^. The study cohort likely represented a higher socioeconomic stratum, as the device and smartphone application are part of a subscription model. Given that our goal was to create and characterize a biometric profile of individuals with regular cycles, we also limited our cohort to participants with median cycle lengths of 21–35 days, excluded cycles shorter than 15 or longer than 45 days, and omitted participants using hormonal contraception, who were pregnant, or who reported perimenopausal symptoms during the data days analyzed. While the within-participant natural experiments offered unique insights into potential causal relationships (Figure 4), the broader population-level and within-participant findings (Figures 2 and 3), remain associative limiting their direct extrapolation to populations with irregular cycles, diverse health conditions, varied socioeconomic backgrounds, or differing ancestries.

Our analyses relied solely on objective biometric and behavioral data, omitting qualitative experiences such as symptom reports or disease status, which could provide crucial context for physiological observations. This absence of qualitative data means the study could not account for subjective experiences or symptomatic presentations. Moreover, additional influence of unmeasured exogenous factors (e.g., environment, socio-cultural elements) and endogenous factors (e.g., stress, illness) could affect the reported outcomes. Our analyses of sleep impacts were constrained to short-term changes (weekly) and we expect that longer term changes in sleep could lead to different physiological responses. Finally, analytical inferences were limited by participant representation for certain conditions (e.g., sleep durations less than 6 hours or equal to/exceeding 9.5 hours; ages greater than 46 years; BMIs less than or equal to 18). The influence of subjective and exogenous factors, as well as longer-term changes and more extreme changes in behavior are all valuable areas for future research on female health and performance using digital health tools.

## Conclusion

This work provides an observational, ecologically valid view of the interplay between sleep and biometrics through the menstrual cycle, and the associations between sleep and menstrual cycle length. Our approach using passive collection of large-scale biometric data and statistical models to understand trends in those data offer significant opportunities for advancing menstrual cycle research and women’s health in general^54^. The longitudinal dataset further revealed the extent and timing of physiological effects associated with sleep changes, a first exploration of behavior and physiology across the menstrual cycle. This study lays the groundwork for prospective intervention studies designed to help individuals better understand and manage physiological fluctuations through the menstrual cycle and elevate their health and athletic performance.

## Acknowledgements

We thank Bill von Hippel and Finn Fielding for sharing their expertise and scientific insight to support this work. We also thank all the participants who shared data for this study. This study was funded by the Wu Tsai Human Performance Alliance at Stanford University and the Joe and Clara Tsai Foundation. The funder played no role in study design, data collection, analysis and interpretation of data, or the writing of this manuscript.

## Author Contributions

AG and JO performed the statistical analyses and generated the figures. AG created the analytical framework. SCJ and JO reviewed pertinent literature. AG, JO, SD, and JH wrote the manuscript. All authors interpreted the data and reviewed the manuscript.

## Methods

### Dataset

We analyzed data from participants who were regular users of a WHOOP device and its associated subscription smartphone application and consented to participate in a women’s health research study. The wearable device (WHOOP 3.0 and WHOOP 4.0) was typically worn on the wrist and passively collected biometric data including nightly measures of resting heart rate (RHR), respiratory rate (RR), heart rate variability (HRV; calculated as root mean square of successive differences), skin temperature (Temp.), and blood oxygen saturation level (Blood O2). Additionally, nightly sleep metrics and daily workout metrics were recorded (see “Daily menstrual, sleep, and biometric data” section for details). The biometrics and sleep measures captured by these devices have been previously validated^55–57^. Participants manually logged their menstruation status, menopause symptoms, and contraception use in the WHOOP smartphone application. Data handling and analysis was approved by and conducted in accordance with the guidelines of the Stanford University Institutional Review Board.

### Inclusion criteria

The full initial dataset comprised 12,010 participants. We analyzed a subset of participants who consistently wore the device, consistently logged menstrual cycles, and experienced regular cycles, as defined below. Participants were 18 to 50 years of age at the time of data collection and were excluded if they reported using hormonal birth control or did not answer the survey questions related to hormonal contraceptive use. Participants who logged pregnancy or menopause symptoms during the data collection period were excluded from the study cohort. Participants were required to demonstrate consistent device wear (>75% of nightly wear) and to have logged at least 90% of their expected cycles based on their median cycle length. A minimum of five recorded menstrual cycles was necessary for inclusion.

In accordance with established guidelines for menstrual cycle research^58^, we only included participants whose median cycle length was between 21 and 35 days. After identifying participants whose median cycle lengths fell in this range, we excluded individual cycles with lengths shorter than 15 days or longer than 45 days. The final number of participants included in the study was 2596 with a total of 42,759 menstrual cycles (Supplementary Table 1).

### Daily menstrual, sleep, and biometric data

For each study participant, we constructed a comprehensive daily time series by integrating menstrual, behavioral, and biometric data. Daily reports of menstrual status were processed to identify cycle start dates. Then we labeled three one-week phases of each menstrual cycle based on cycle start dates: premenstrual (7 days prior to cycle onset through 1 day prior), menstrual (day of cycle onset and the subsequent 6 days), and postmenstrual (7 to 14 days post cycle onset). Cycle days outside of these weeks were labeled “other”.

Daily sleep variables included sleep duration and sleep onset time. We computed and included daily workout data summary metrics: duration, timing, and time in heart rate zones (5 zones as a function of max heart rate, z1= 50-59%, z2=60-69%, z3=70-79%, z4=80-89%, z5=90-100%). Max heart rate was either provided by the user through the app or computed using Gellish’s formula^59^. We computed overall daily workout intensity with Edward’s TRIMP (eTRIMP, Edwards, S., (1993) Heart Rate Monitor Book. Polar CIC Inc.), which is heart rate zone times duration in minutes (e.g., eTRIMP=60 for 60 minutes in HR zone 1 or 12 minutes in zone 5).

Nightly biometric data (RHR, HRV, RR, Temp., Blood O2) were included and further processed (see Biometrics signal processing section). Not all timepoints contained the five biometrics: 2,114 participants switched from WHOOP 3.0 devices (collecting RHR, HRV and RR) to WHOOP 4.0 devices (additionally collecting Temp. and Blood O2). This resulted in 15% of the data without Temp. and Blood O2 measures. Across participants, we analyzed a total of 1,298,555 days of data (Supplementary Table 1).

### Biometrics signal processing

After constructing the time-series data of daily biometrics, we interpolated, filtered, and normalized each individual’s daily biometric values. We used linear interpolation to fill in missing daily biometric data for gaps of 7 days or fewer in length, with longer gaps imputed to the participant’s mean value for that biometric (temporarily during filtering and normalization). We then processed the daily biometric measurements using a bandpass Butterworth filter (2nd order) with frequency bounds designed to remove long-term drift (high-pass: 1/90 days−1) and short-term noise (low-pass: 1/10 days−1). We applied phase correction to the filtered signal by temporally shifting it backwards by half the filter order to account for the processing delay.

Normalized daily biometrics for each participant were then defined as the percentage change from an individual’s average (filtered signal /signal average x 100) (Figure S7). In order to preserve missing data patterns in the data, the previously mean-imputed data days were replaced with NaN values. Normalizing the data enabled us to compare physiological patterns across individuals with different baseline values of their biometrics. We report the across-participant average biometric value for varying ages, BMIs, and sleep durations (Figure S4, S5, S10) to give a sense for the magnitude of the raw biometric signals across these different groups.

### Cycle-level menstrual and sleep data

*Cycle length* was computed as the duration in days between two consecutive cycle start dates. *Cycle length deviation* was defined as a binary variable: “True” if the cycle length was different from that participant’s median cycle length by three days or more, and “False” otherwise. The three-day threshold for deviation was chosen to account for variability in participants logging their cycles and prior research showing that minor cycle length deviations are expected.

Average sleep duration for a cycle was computed from the nightly sleep durations auto-detected by the WHOOP device^55,56^. Sleep duration variability was defined as the variance in sleep duration over a cycle. When used in statistical analyses, we converted the variance into a logarithmic scale as a variance stabilization procedure. When plotting sleep duration variability (Figure 2), we used standard deviations for interpretability. Sleep duration variability bins were selected to be linear in the logarithmic scale, corresponding to standard deviation bin edges of [15, 30, 42, 60, 85, 120]. Circular variance was used to compute sleep onset variability, while the cosine and sine components of sleep onset time were averaged by cycle.

### Hypothesis testing and inference

Statistical results in this work are the product of a Generalized Estimating Equation framework, with individual menstrual cycles as the observations and an exchangeable covariance structure per participant. The covariates and modeling details varied depending on the specific analysis and outcome of interest (Supplementary Table 3). All covariates and predictors in the statistical analyses were continuous. Regression weights were computed based on the binned cycle counts of the predictor variable to obtain regression coefficients that appropriately represent the span of the predictor. These weights were computed as 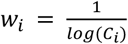, where *C*_*i*_ is the total number of observations of group “*i*”.

After model fitting, we computed the conditional expectation of the outcome at different values of the predictor of interest while holding the other variables at the “average participant” level. For example, for the effects of sleep duration on the mean cycle shown in Figure 2a, the regression line and CI were the result of varying sleep duration as a model input while keeping the other covariates at their mean values (e.g., age=33 and BMI=25).

All models included age and BMI, along with their squared terms as covariates or predictors of interest. Yearly seasonality effects were accounted for by taking the cosine and sine component of the cycle onset day^27^. Models that predicted cycle length deviations had median cycle length as a covariate. Sleep covariates or predictors included sleep duration and its square, sleep duration variability and its square, sleep onset, and sleep onset variability. Workout covariates included workout duration, eTRIMP and its square, workout mean intensity (eTRIMP/duration), and average workout onset. Sleep and workout onset times were decomposed into cosine and sine components to address their circularity. In the statistical models that quantified changes in behavior, the chronic term was the 3-week average up to the point of change (e.g., the 21-day average sleep before a menstrual week).

### Biometric modeling

We fit Generalized Additive Models (GAM) using Bayesian Additive Models to disentangle the effects of age and cycle length on physiological variables throughout the menstrual cycle. For each biometric, data were aggregated by participant and cycle length, spanning 10 days prior to 35 days post cycle onset. Daily sleep duration and exercise training impulse (eTRIMP), along with their 21-day moving averages, were included as behavioral covariates, with exercise metrics lagged by one day to align with physiological responses. Additional confounders included BMI, seasonality (cosine and sine terms for yearly periodicity), and weekend effects (modeled as a binary, “True” Saturday and Sunday, “False” otherwise).

Age-cycle length interactions were modeled through smooth terms for cycle day, cycle length, and age, with tensor product interactions capturing how biometric trajectories vary jointly with age and cycle length. Behavioral covariates were similarly modeled with smooth terms and tensor interactions with cycle day. Additional interaction tensor products were included for sleep duration and eTRIMP. Models were fitted using an AR(1) correlation structure to account for temporal autocorrelation within subject-cycle combinations, employing fast restricted maximum likelihood estimation with automatic smoothness selection (R package MGCV).

To visualize the primary effects of age and cycle length, simulation data were generated with behavioral covariates fixed at population averages (sleep duration = 7.25 hours, eTRIMP = 80, BMI=25). Model performance was evaluated using deviance explained (D^2^) and effective degrees of freedom.

The temporal relationship between biometrics was quantified using a Vector Autoregression model with a lag of 3 days (Figure S7). This means that the model can predict a biometric at time step *t+1* with a linear combination of all the biometrics at time *t* through time step *t-3*. For this analysis, we processed the raw data with a low-pass filter of 1/3 days^-1^ to allow for faster fluctuations within biometrics and capture the dynamical structure between biometrics (as opposed to the 1/10 days^-1^ in all other analyses). Other processing steps were as in the main analysis (imputation, high-pass filtering, and normalization). The resulting normalized biometrics were then fed into Python’s *VAR* function in the *statsmodels* package.

### Changes in sleep and biometrics across the cycle

We quantified the likelihood of changing sleep duration during a given cycle week (premenstrual, menstrual, postmenstrual; Figure 4a). We selected cycle-weeks for which sleep duration changes were preceded by periods of stable sleep (three-week average between 7-8 hours). This approach captured changes from chronic behavior while minimizing overlap with the cycle week of interest (e.g., data from the 10th menstrual week would not overlap with data from the 9th menstrual week in a 28-day cycle). We binned sleep duration percentage changes as follows: [-25, 10], (−10, −5], (−5, 5], (5, 10], (10, 25] across menstrual weeks. A 10% change from a 7-hour average sleep corresponded to a 42-minute change in sleep duration for that week. These bins then represented meaningful changes in sleep behavior, and their frequencies were used as regression weights in the subsequent biometric analyses (see Hypothesis testing and inference section). We labeled the continuous biometrics according to the binned weekly behavior changes for plotting purposes. Biometric changes were modeled with GEEs for the last day of the week in which we observed changes in sleep duration (day −1 for premenstrual, day 6 for menstrual, day 13 for postmenstrual; dots in Figure 4b,c).

## Code availability

Python code will be made available at the time of publication on a GitHub repository. The underlying code was written in Python (version 3.11) and R (version 4.3.3).

## Data availability

The data that support the findings of this study are available from WHOOP, Inc., but restrictions apply to the availability of these data and so they are not publicly available. Data are however available from the authors upon reasonable request and with permission of WHOOP, Inc. Statistical weights from GAM analyses will be made available at the time of publication on a GitHub repository.

## References

1. Nowakowski, S., Meers, J. & Heimbach, E. Sleep and women’s health. Sleep Med. Res. 4, 1–22 (2013).

2. Rugvedh, P., Gundreddy, P. & Wandile, B. The Menstrual Cycle’s Influence on Sleep Duration and Cardiovascular Health: A Comprehensive Review. Cureus 15, 10 (2023).

3. Carmichael, M. A., Thomson, R. L., Moran, L. J. & Wycherley, T. P. The Impact of Menstrual Cycle Phase on Athletes’ Performance: A Narrative Review. Int. J. Environ. Res. Public Health 18, 1667 (2021).

4. Rosen Vollmar, A. K., Mahalingaiah, S. & Jukic, A. M. The menstrual cycle as a vital sign: a comprehensive review. F S Rev. 6, 100081 (2025).

5. Li, K. et al. Characterizing physiological and symptomatic variation in menstrual cycles using self-tracked mobile-health data. NPJ Digit. Med. 3, 79 (2020).

6. Stellingwerff, T. et al. Review of the scientific rationale, development and validation of the International Olympic Committee Relative Energy Deficiency in Sport Clinical Assessment Tool: V.2 (IOC REDs CAT2)—by a subgroup of the IOC consensus on REDs. Br. J. Sports Med. 57, 1109–1121 (2023).

7. Baker, F. C., Siboza, F. & Fuller, A. Temperature regulation in women: Effects of the menstrual cycle. Temperature (Austin) 7, 226–262 (2020).

8. Lin, G. et al. Understanding wrist skin temperature changes to hormone variations across the menstrual cycle. NPJ Womens Health 2, 35 (2024).

9. Marshall, J. A field trial of the basal-body-temperature method of regulating births. Lancet 292, 8–10 (1968).

10. Shilaih, M., Clerck, V. D., Falco, L., Kübler, F. & Leeners, B. Pulse Rate Measurement During Sleep Using Wearable Sensors, and its Correlation with the Menstrual Cycle Phases, A Prospective Observational Study. Sci. Rep. 7, 1294 (2017).

11. Moran, V. H., Leathard, H. L. & Coley, J. Cardiovascular functioning during the menstrual cycle. Clin. Physiol. 20, 496–504 (2000).

12. Alzueta, E. et al. Tracking Sleep, Temperature, Heart Rate, and Daily Symptoms Across the Menstrual Cycle with the Oura Ring in Healthy Women. Int. J. Womens Health 14, 491–503 (2022).

13. Tenan, M. S., Brothers, R. M., Tweedell, A. J., Hackney, A. C. & Griffin, L. Changes in resting heart rate variability across the menstrual cycle. Psychophysiology 51, 996–1004 (2014).

14. Sims, S. T., Ware, L. & Capodilupo, E. R. Patterns of endogenous and exogenous ovarian hormone modulation on recovery metrics across the menstrual cycle. BMJ Open Sport Exerc. Med. 7, e001047 (2021).

15. Schmalenberger, K. M. et al. A Systematic Review and Meta-Analysis of Within-Person Changes in Cardiac Vagal Activity across the Menstrual Cycle: Implications for Female Health and Future Studies. J. Clin. Med. 8, 1946 (2019).

16. Jasinski, S. R., Presby, D. M., Grosicki, G. J., Capodilupo, E. R. & Lee, V. H. A Novel method for quantifying fluctuations in wearable derived daily cardiovascular parameters across the menstrual cycle. NPJ Digit Med 7, 373 (2024).

17. Keeler Bruce, L., González, D., Dasgupta, S. & Smarr, B. L. Biometrics of complete human pregnancy recorded by wearable devices. NPJ Digit. Med. 7, 207 (2024).

18. Jasinski, S. R., Rowan, S., Presby, D. M., Claydon, E. A. & Capodilupo, E. R. Wearable-derived maternal heart rate variability as a novel digital biomarker of preterm birth. PLoS One 19, e0295899 (2024).

19. Arey, L. B. The Degree of Normal Menstrual Irregularity**Contribution No. 245. Acknowledgment is due several persons who have supplied information to further the completion of this report. Drs. E. Allen, C. F. Fluhmann, J. L. King, L. J. Latz, J. P. Pratt, and M. C. Shelesnyak have cooperated by sending miscellaneous, supplementary data to clarify and extend their published accounts of group studies. Am. J. Obstet. Gynecol. 37, 12–29 (1939).

20. Chiazze, L., Jr, Brayer, F. T., Macisco, J. J., Jr, Parker, M. P. & Duffy, B. J. The Length and Variability of the Human Menstrual Cycle. JAMA 203, 377–380 (1968).

21. Gunn, D. L., Jenkin, P. M. & Gunn, A. L. Menstrual Periodicity; Statistical Observations on a Large Sample of Normal Cases. BJOG 44, 839–879 (1937).

22. Lenton, E. A., Landgren, B., Sexton, L. & Harper, R. Normal variation in the length of the follicular phase of the menstrual cycle: effect of chronological age. BJOG 91, 681–684 (1984).

23. Bull, J. R. et al. Real-world menstrual cycle characteristics of more than 600,000 menstrual cycles. NPJ Digit. Med. 2, 83 (2019).

24. Cunningham, A. C. et al. Chronicling menstrual cycle patterns across the reproductive lifespan with real-world data. Sci. Rep. 14, 10172 (2024).

25. Li, H. et al. Menstrual cycle length variation by demographic characteristics from the Apple Women’s Health Study. NPJ Digit. Med. 6, 100 (2023).

26. Goodale, B. M. et al. Wearable Sensors Reveal Menses-Driven Changes in Physiology and Enable Prediction of the Fertile Window: Observational Study. J Med Internet Res 21, e13404 (2019).

27. Pierson, E., Althoff, T., Thomas, D., Hillard, P. & Leskovec, J. Daily, weekly, seasonal and menstrual cycles in women’s mood, behaviour and vital signs. Nat. Hum. Behav. 5, 716–725 (2021).

28. Baker, F. C. & Driver, H. S. Circadian rhythms, sleep, and the menstrual cycle. Sleep Med. 8, 613–622 (2007).

29. Alzueta, E. & Baker, F. C. The Menstrual Cycle and Sleep. Sleep Med. Clin. 18, 399–413 (2023).

30. McNulty, K. L. et al. The Effects of Menstrual Cycle Phase on Exercise Performance in Eumenorrheic Women: A Systematic Review and Meta-Analysis. Sports Med. 50, 1813–1827 (2020).

31. Reed, B. G. & Carr, B. R. The normal menstrual cycle and the control of ovulation. (2015).

32. Liu, Y., Gold, E. B., Lasley, B. L. & Johnson, W. O. Factors Affecting Menstrual Cycle Characteristics. Am. J. Epidemiol. 160, 131–140 (2004).

33. Sletten, T. L. et al. The importance of sleep regularity: a consensus statement of the National Sleep Foundation sleep timing and variability panel. Sleep Health 9, 801–820 (2023).

34. Umetani, K., Singer, D. H., McCraty, R. & Atkinson, M. Twenty-Four Hour Time Domain Heart Rate Variability and Heart Rate: Relations to Age and Gender Over Nine Decades. J. Am. Coll. Cardiol. 31, 593–601 (1998).

35. Hickcox, L., Bates, S., Hashemzadeh, M. & Movahed, M. R. Higher Heart Rate Is Independently Associated With Abnormal Body Mass Index in a J Shape Pattern. Crit. Pathw. Cardiol. 22, (2023).

36. Van Reen, E. & Kiesner, J. Individual differences in self-reported difficulty sleeping across the menstrual cycle. Arch. Womens. Ment. Health 19, 599–608 (2016).

37. Hachul, H. et al. Does the reproductive cycle influence sleep patterns in women with sleep complaints? Climacteric 13, 594–603 (2010).

38. Kennedy, K. E. R. et al. Menstrual regularity and bleeding is associated with sleep duration, sleep quality and fatigue in a community sample. J. Sleep Res. 31, e13434 (2022).

39. Lim, A. J. R., Huang, Z., Chua, S. E., Kramer, M. S. & Yong, E.-L. Sleep Duration, Exercise, Shift Work and Polycystic Ovarian Syndrome-Related Outcomes in a Healthy Population: A Cross-Sectional Study. PLoS One 11, e0167048 (2016).

40. Komada, Y. et al. Social jetlag and menstrual symptoms among female university students. Chronobiol. Int. 36, 258–264 (2019).

41. Labyak, S., Lava, S., Turek, F. & Zee, P. Effects of shiftwork on sleep and menstrual function in nurses. Health Care Women Int. 23, 703–714 (2002).

42. Lateef, O. M. & Akintubosun, M. O. Sleep and Reproductive Health. J. Circadian Rhythms18, 1 (2020).

43. Yu, J.-L. et al. Tracking of menstrual cycles and prediction of the fertile window via measurements of basal body temperature and heart rate as well as machine-learning algorithms. Reprod. Biol. Endocrinol. 20, 118 (2022).

44. Su, H.-W., Yi, Y.-C., Wei, T.-Y., Chang, T.-C. & Cheng, C.-M. Detection of ovulation, a review of currently available methods. Bioeng. Transl. Med. 2, 238–246 (2017).

45. Bauman, J. E. Basal body temperature: unreliable method of ovulation detection. Fertil. Steril. 36, 729–733 (1981).

46. Wang, Y. et al. Performance of algorithms using wrist temperature for retrospective ovulation day estimate and next menses start day prediction: a prospective cohort study. Hum. Reprod. 40, 469–478 (2025).

47. Saleh, T. M. & Connell, B. J. 17β-Estradiol modulates baroreflex sensitivity and autonomic tone of female rats. J. Auton. Nerv. Syst. 80, 148–161 (2000).

48. Brown, A. M. C. & Gervais, N. J. Role of Ovarian Hormones in the Modulation of Sleep in Females Across the Adult Lifespan. Endocrinology 161, bqaa128 (2020).

49. Dettoni, J. L. et al. Cardiovascular effects of partial sleep deprivation in healthy volunteers. J. Appl. Physiol. 113, 232–236 (2012).

50. Meier-Ewert, H. K. et al. Effect of sleep loss on C-Reactive protein, an inflammatory marker of cardiovascular risk. J. Am. Coll. Cardiol. 43, 678–683 (2004).

51. Kaciuba-Uscilko, H. & Grucza, R. Gender differences in thermoregulation: Curr. Opin. Clin. Nutr. Metab. Care 4, 533–536 (2001).

52. Pfender, E. J., Kuijpers, K. L., Wanzer, C. V. & Bleakley, A. Cycle Syncing and TikTok’s Digital Landscape: A Reasoned Action Elicitation Through a Critical Feminist Lens. Qual. Health Res. 10497323241297683 (2024).

53. Mikkonen, R. S. & Häkkinen, K. Evidence for Periodizing Strength and/or Endurance Training According to Menstrual Cycle Phases to Optimize Female Athlete Performance Is Lacking. Strength Cond. J. (2025).

54. Turco, M. Y. & Kraft, O. A holistic approach to advancing women’s health. Nat. Rev. Bioeng. 3, 432–434 (2025).

55. Berryhill, S. et al. Effect of wearables on sleep in healthy individuals: a randomized crossover trial and validation study. J. Clin. Sleep Med. 16, 775–783 (2020).

56. Miller, D. J. et al. A Validation Study of a Commercial Wearable Device to Automatically Detect and Estimate Sleep. Biosensors (Basel) 11, (2021).

57. Miller, D. J., Sargent, C. & Roach, G. D. A Validation of Six Wearable Devices for Estimating Sleep, Heart Rate and Heart Rate Variability in Healthy Adults. Sensors 22, (2022).

58. Schmalenberger, K. M. et al. How to study the menstrual cycle: Practical tools and recommendations. Psychoneuroendocrinology 123, 104895 (2021).

59. Gellish, R. L. et al. Longitudinal modeling of the relationship between age and maximal heart rate. Med. Sci. Sports Exerc. 39, 822–829 (2007).

