## Supplementary Information for "The menstrual cycle through the lens of a wearable device: insights into physiology, sleep, and cycle variability"

#### Supplementary Tables

**Supplementary Table 1. Participant Data and Demographics**

| | |
|---|---|
| Participants (#) | 2,596 |
| Data days (#) | 1,298,555 |
| Data days with cardiorespiratory biometrics (#) | 1,126,211 |
| Data days with all biometrics (#) | 914,440 |
| Journaled menstrual cycles (#) | 42,759 |
| | Across Participants Mean ± S.D. |
| Age (years) | 33.7 ± 6.5 |
| Body mass index (BMI) | 24.8 ± 4.4 |
| Journaled menstrual cycles (#) | 18.2 ± 4.5 |
| Expected cycles (#) | 18.0 ± 4.5 |
| Data days (#) | 500 ± 116 |
| Sleep duration (hours) | 7.2 ± 0.6 |
| Workout duration (mins/day) | 40 ± 31 |

**Supplementary Table 2. Within-Cycle Biometrics**

| | Mean | Standard Deviation | Within-Cycle Range | Within-Cycle Range [%] |
|---|---|---|---|---|
| Resting heart rate [beats/min] | 60.2 ± 7.5 | 2.4 ± 0.6 | 8.1 ± 2.2 | 13.7 ± 3.5 |
| Heart rate variability [ms] | 59.3 ± 25.2 | 6.4 ± 3.3 | 22.0 ± 11.2 | 37.8 ± 12.4 |
| Respiratory rate [breaths/min] | 15.9 ± 1.5 | 0.3 ± 0.1 | 1.0 ± .0.2 | 6.4 ± 1.6 |
| Skin temperature [degrees C] | 34.3 ± 0.6 | 0.3 ± 0.1 | 0.9 ± 0.2 | 2.8 ± 0.7 |
| Blood oxygen saturation level [%] | 95.8 ± 1.2 | 0.6 ± 0.2 | 2.1 ± 0.7 | 2.2 ± 0.7 |

For each participant and biometric, we compute the by-cycle mean, standard deviation, and the range (peak to trough difference), then compute the across-cycle averages, and report the resulting mean and standard deviation across participants.

#### Supplementary Table 3. GEE Statistical Analyses Summary

| | Outcome | Predictor(s) of interest | Covariates: (age, BMI, and season included in all models) | Sample Size C: #cycles, P: #participants | Fit Family |
|---|---|---|---|---|---|
| Fig. 1a | Cycle length | age | | C=42,759; P=2,596 | Gaussian |
| Fig. 1b | Cycle length deviation | age | median cycle length, age x median cycle length | C=42,759; P=2,596 | Binomial |
| Fig. 2a | Cycle length | sleep duration | sleep metrics, workout metrics | C=39,079; P=2,595 | Gaussian |
| Fig. 2b | Cycle length deviation | sleep duration | median cycle length, age x median cycle length, sleep metrics, workout metrics | C=39,079; P=2,595 | Binomial |
| Fig. 2c | Cycle length | sleep variability | sleep metrics, workout metrics | C=39,079; P=2,595 | Gaussian |
| Fig. 2d | Cycle length deviation | sleep variability | median cycle length, age x median cycle length, sleep metrics, workout metrics | C=39,079; P=2,595 | Binomial |
| Fig. 4a | 10% change in sleep | menstrual cycle week (MCW) | cycle length, sleep metrics*, workout metrics, MCW x chronic sleep | C=20,814; P=2,312 | Binomial |
| Fig. 4c, Fig S11 | Percent change in RHR** or another biometric | percent change in sleep duration (CIS) | cycle length, MCW, chronic sleep, sleep metrics*, workout metrics, MCW x (CIS, chronic sleep, cycle length) age x cycle length, CIS x chronic sleep | Cardiorespiratory: C=20,806; P=2,312<br>Temp. and Blood O2 C=17,917; P=2,228 | Gaussian |
| Fig S9b | Biometric magnitude range | cycle length and age | sleep metrics, workout metrics | Cardiorespiratory: C=39,072; P=2,595<br>Temp. and Blood O2 C=33,560; P=2,595 | Gaussian |

GEE=Generalized Estimating Equations for population level inferences
* sleep duration excluded as it is co-linear with change in sleep when chronic sleep is present
** percent change in resting heart rate at the last day of the menstrual cycle of week (1 day prior to cycle start for premenstrual, day 6 for menstrual, day 13 for postmenstrual) relative to participant's average
MCW = menstrual cycle week, CIS = change in sleep

### Supplementary Figures:

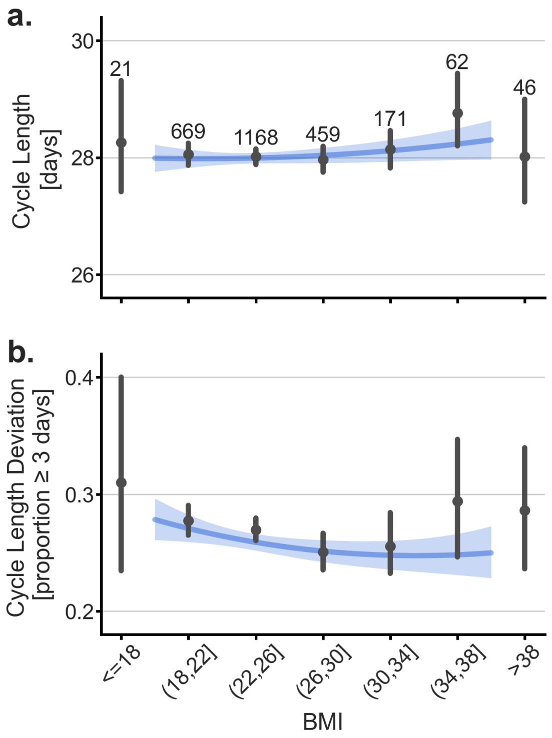


**Figure S1. Cycle length and cycle length variability by body mass index (BMI).**
**(a)** Population mean cycle length (in days) was not different by BMI. We did detect a significant association between BMI and cycle length. **(b)** Cycle length deviation slightly decreased with BMI. Using a GEE model, with age and median cycle length as covariates, we found that cycle length deviation was 0.27 [95%CI: 0.26 - 0.28] at a BMI of 20, and 0.25 [95%CI: 0.24 - 0.26] at a BMI of 30, for an OR of 1.1 [95%CI: 1.0 - 1.2]. Note that 89% of the cohort had a BMI of less than 30 kg/m2, thus we do not make inferences for higher BMIs. BMI (kg/m2) was calculated as the body mass divided by height squared, and grouped into 4-unit bins.

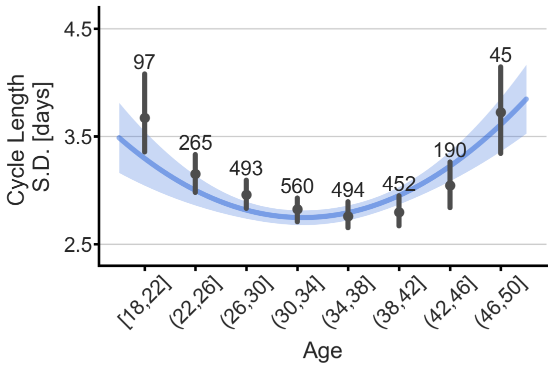


**Figure S2. Cycle length standard deviation (S.D.) by age.** Cycle length standard deviation follows a U-shaped pattern with age. Using a statistical model with cycle length standard deviation as the output we determined that the minimum of deviation was at age 32 [95%CI: 30.6 - 33.7] at 2.7 days, and there were more variable cycle lengths in the early 20's and late 40's (at age 24 it was 2.9, at age 44 it was 3.2).

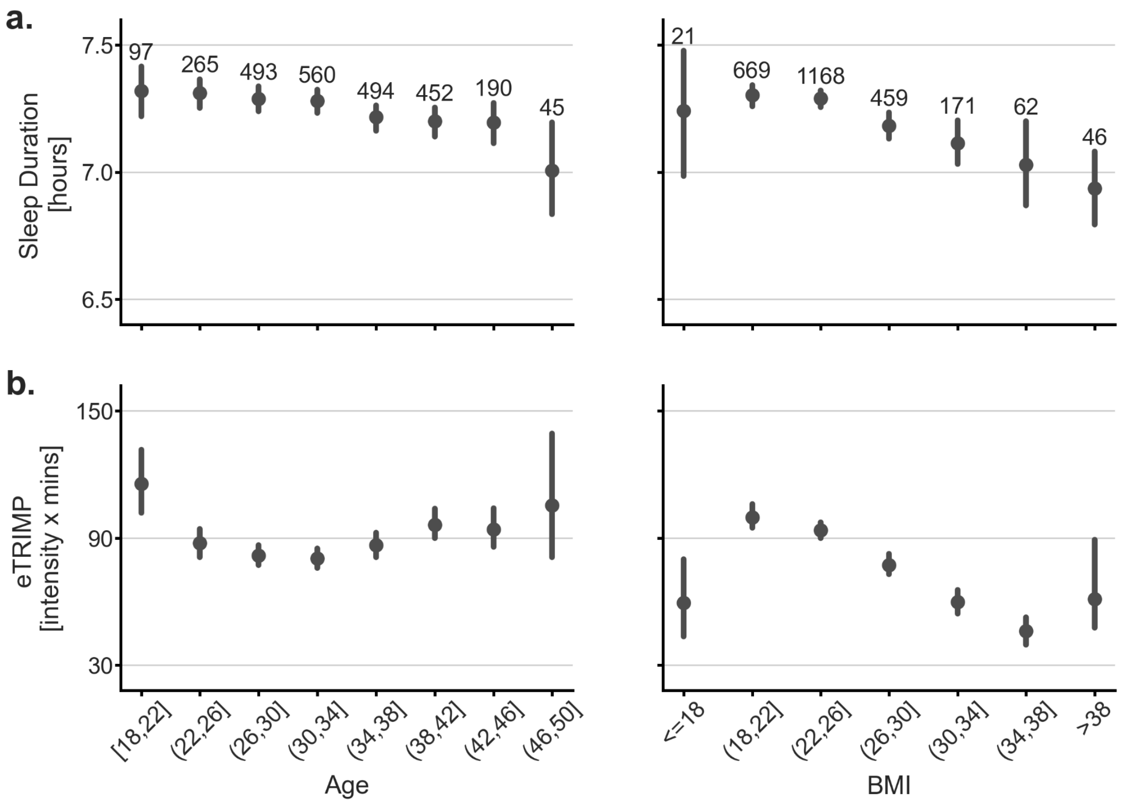


**Figure S3. Sleep duration and workout loads by age and BMI.** **(a)** Average sleep duration decreased with age and BMI. Each point represents the mean sleep duration across participants (left panel) for a given 4-year age bin from 18 - 50 years or BMI bin (18 or lower, 18 to 38 in 4-unit increments, 38 and above). Annotations above the points indicate the number of participants per bin. **(b)** Average workout load was consistent across ages and decreased with BMI. Workout load was computed with Edward's TRIMP (eTRIMP, ref), as a measure of intensity or heart rate zone times duration in minutes (e.g., eTRIMP=60: for 60 HR zone 1 minutes and 12 zone 5 minutes).

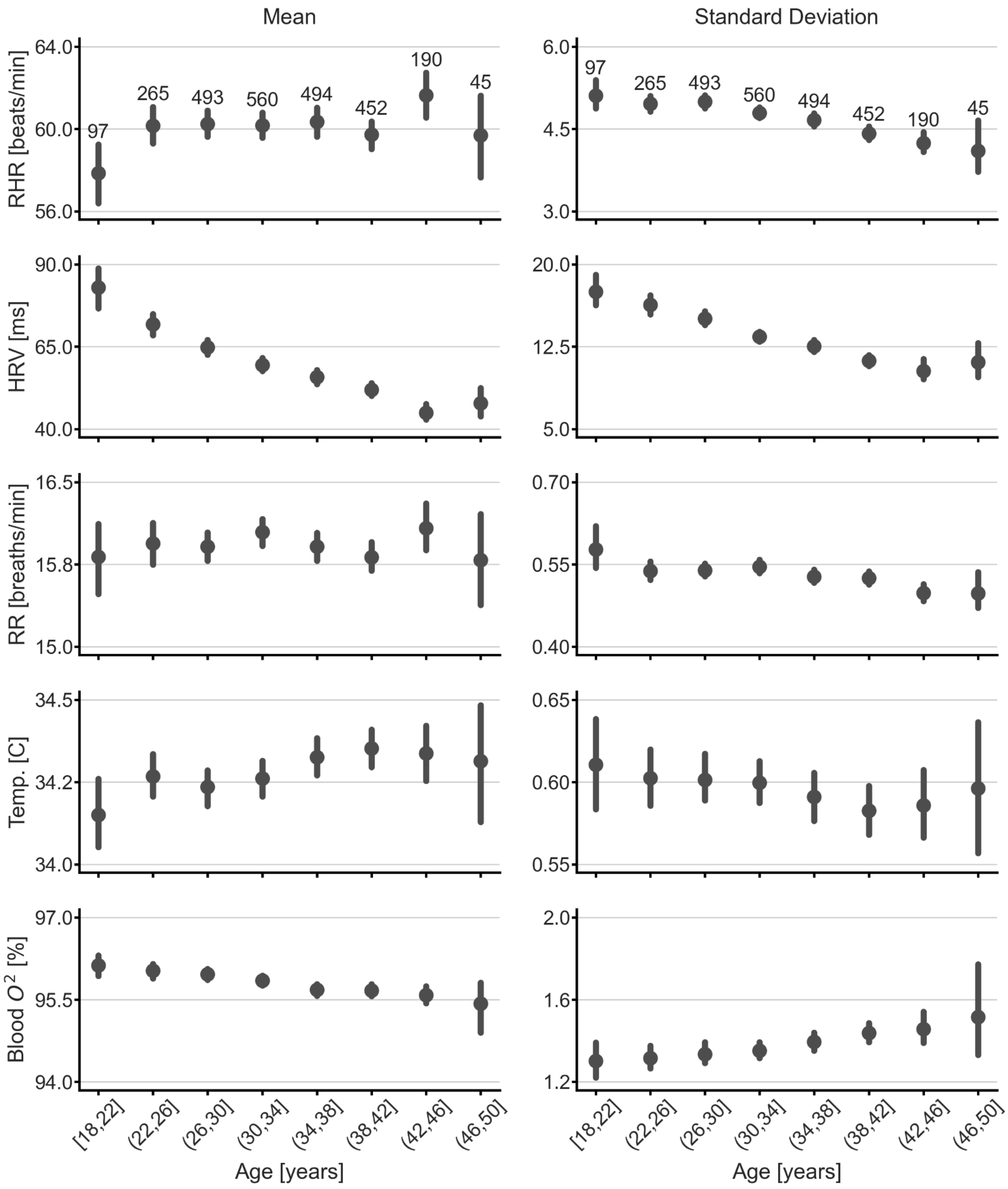


**Figure S4. Raw physiological biometrics by age.** Resting heart rate (RHR), heart rate variability (HRV), respiratory rate (RR), skin temperature (Temp.), and blood oxygen saturation level (Blood O2) levels across ages. Each point represents the (left column) mean value or (right column) standard deviation of that biometric across participants for a given 4-year age bin from 18 - 50 years (x-axis). Annotations above the points indicate the number of participants per bin.

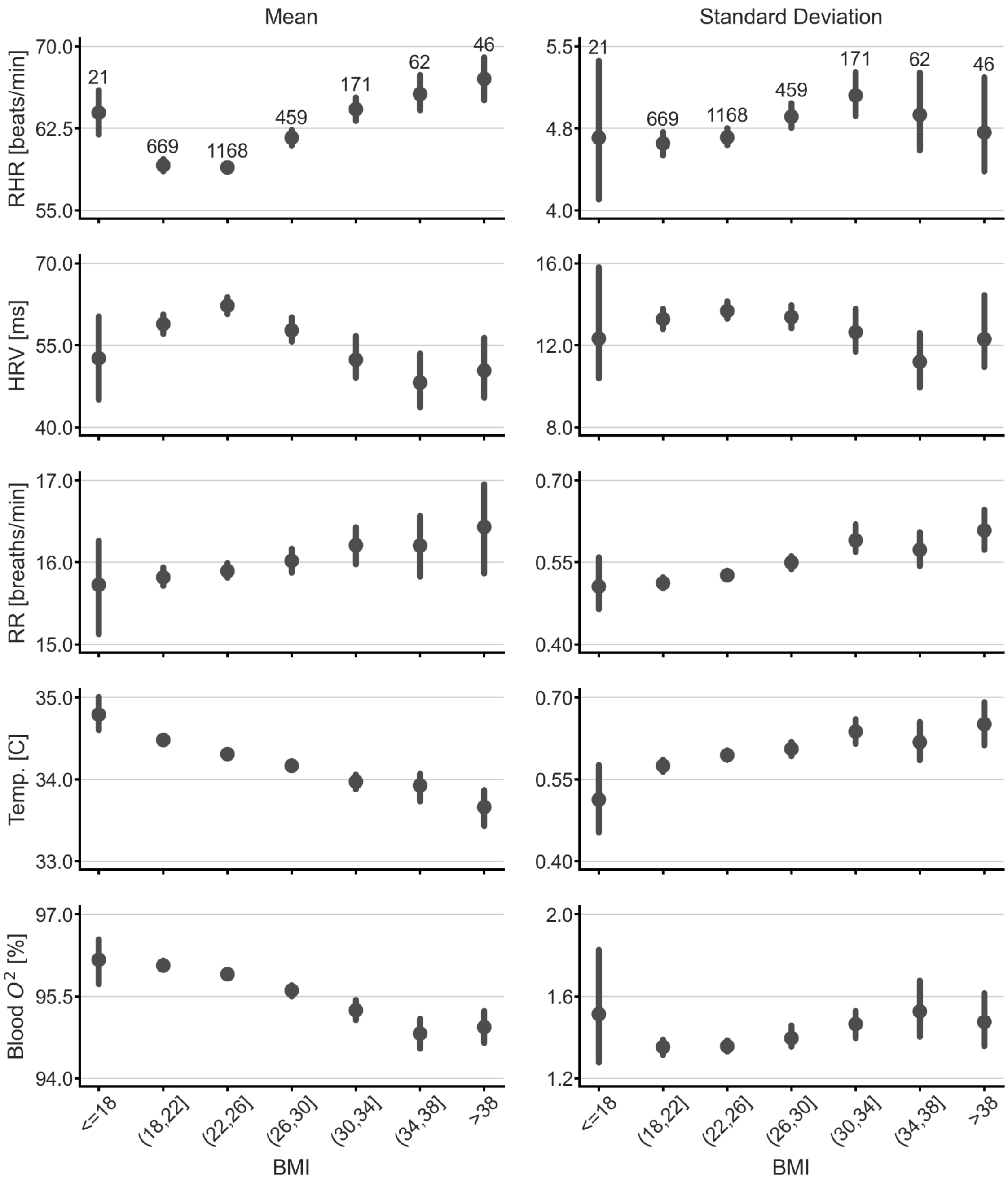


**Figure S5. Raw physiological biometrics by BMI.** Resting heart rate (RHR), heart rate variability (HRV), respiratory rate (RR), skin temperature (Temp.), and blood oxygen saturation level (Blood O2) levels across body mass index (BMI). Each point represents the (left column) mean value or (right column) standard deviation of that biometric across participants for a given BMI bin (x-axis: 18 or lower, 18 to 38 in 4-unit increments, 38 and above). Annotations above the points indicate the number of participants per bin.

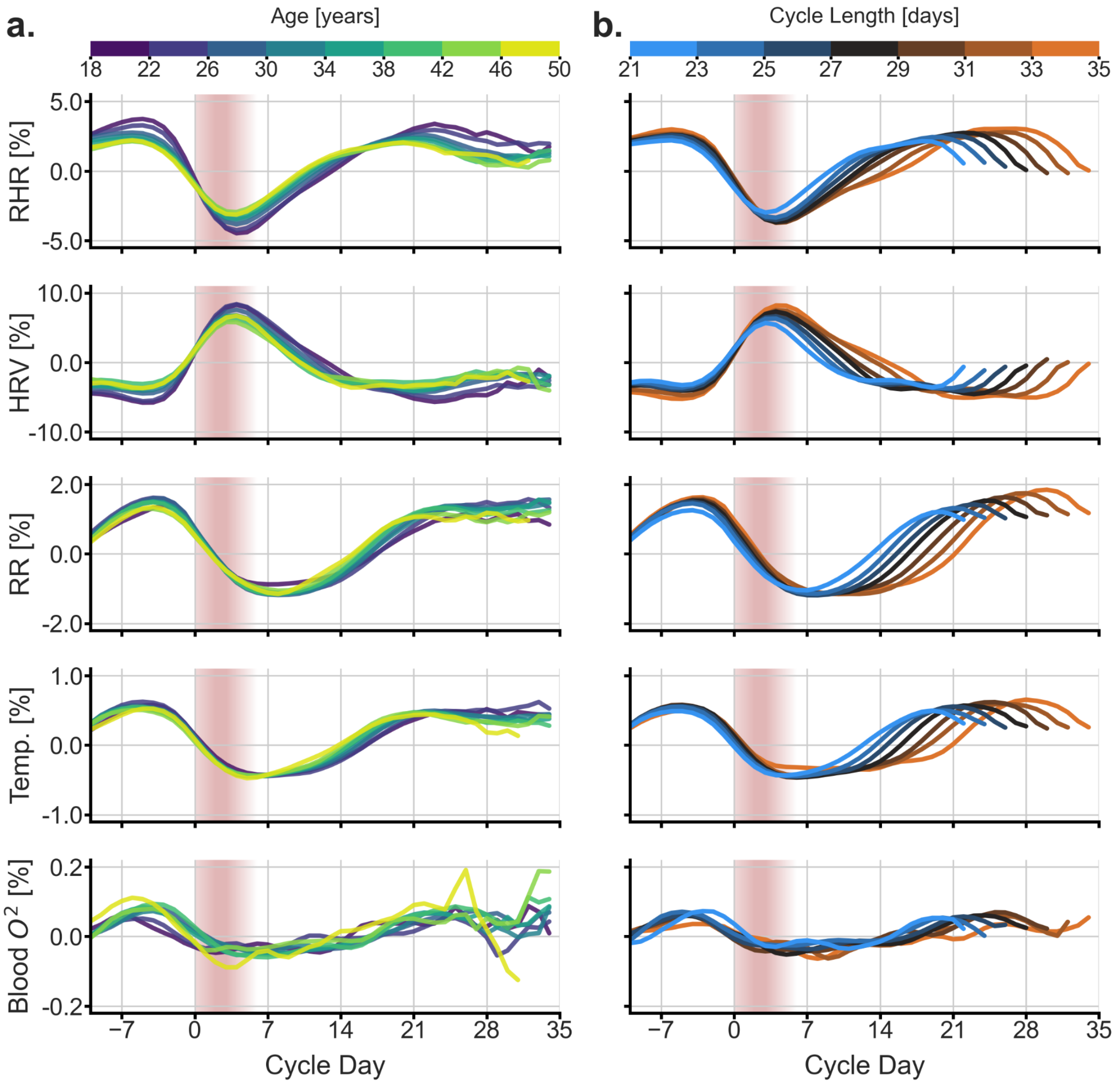


**Figure S6. Normalized biometrics through the menstrual cycle across ages and cycle lengths.** Variation in average nightly physiological biometrics, including resting heart rate (RHR), heart rate variability (HRV), respiratory rate (RR), skin temperature (Temp.), and blood oxygen saturation level (Blood $O^2$) are shown as a function of menstrual cycle day. **(a)** Each colored line represents the average by age group, with participants grouped in 4-year age bins. **(b)** Each colored line represents the biometrics' average across participants that had cycles in that 2-day cycle length bin. Cycle day zero indicates the day of self-reported menstrual cycle onset. Values were z-scored by participant to highlight the magnitude of fluctuations. Curves are based directly on the cohort data and thus age and cycle-length effects can be interacting, in contrast to the model generated curves in Figure 3.

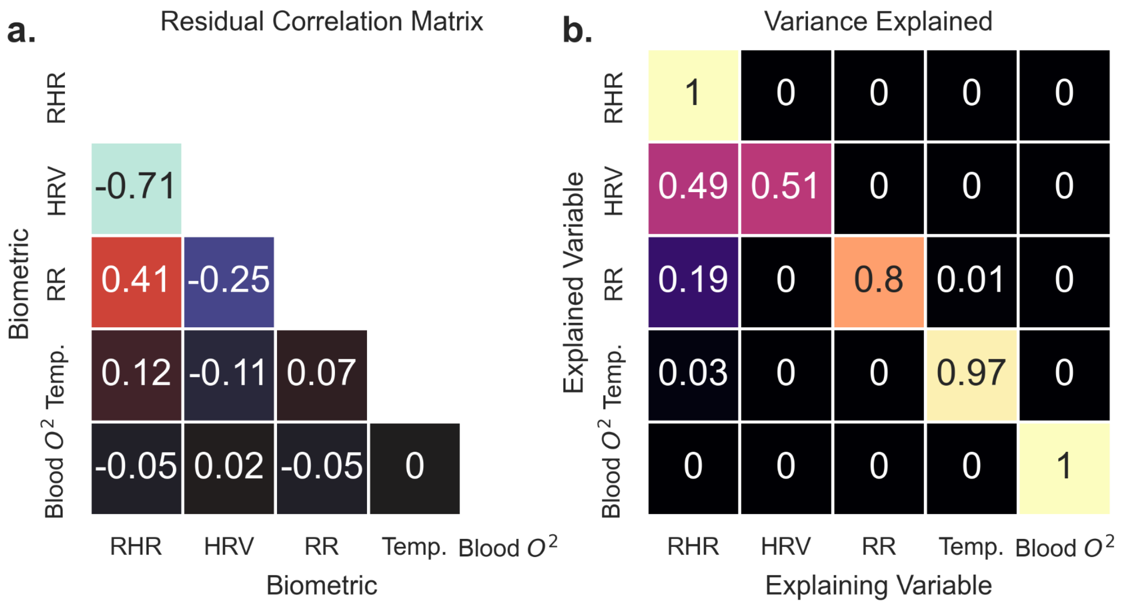


**Figure S7. Relationships between biometrics during the menstrual cycle.**
**(a)** Correlation matrix showing how biometrics change together after removing time-delay effects. Values show correlations between model residuals, indicating which biometrics move up and down together at the same time, beyond what can be predicted from their own past values. The lower triangular mask highlights the symmetric relationships. (**b)** Variance decomposition matrix showing how much each variable's prediction errors are explained by unexpected changes in other variables (3 days ahead). Rows show the variable being predicted, columns show the variable providing the explanation. Diagonal elements show how much a variable explains its own prediction errors, while off-diagonal elements show cross-variable influences. The analysis reveals that heart rate measures (RHR and HRV) were tightly linked, while RHR also influenced breathing rate, consistent with coordinated autonomic nervous system control during the menstrual cycle. See Methods for processing details.

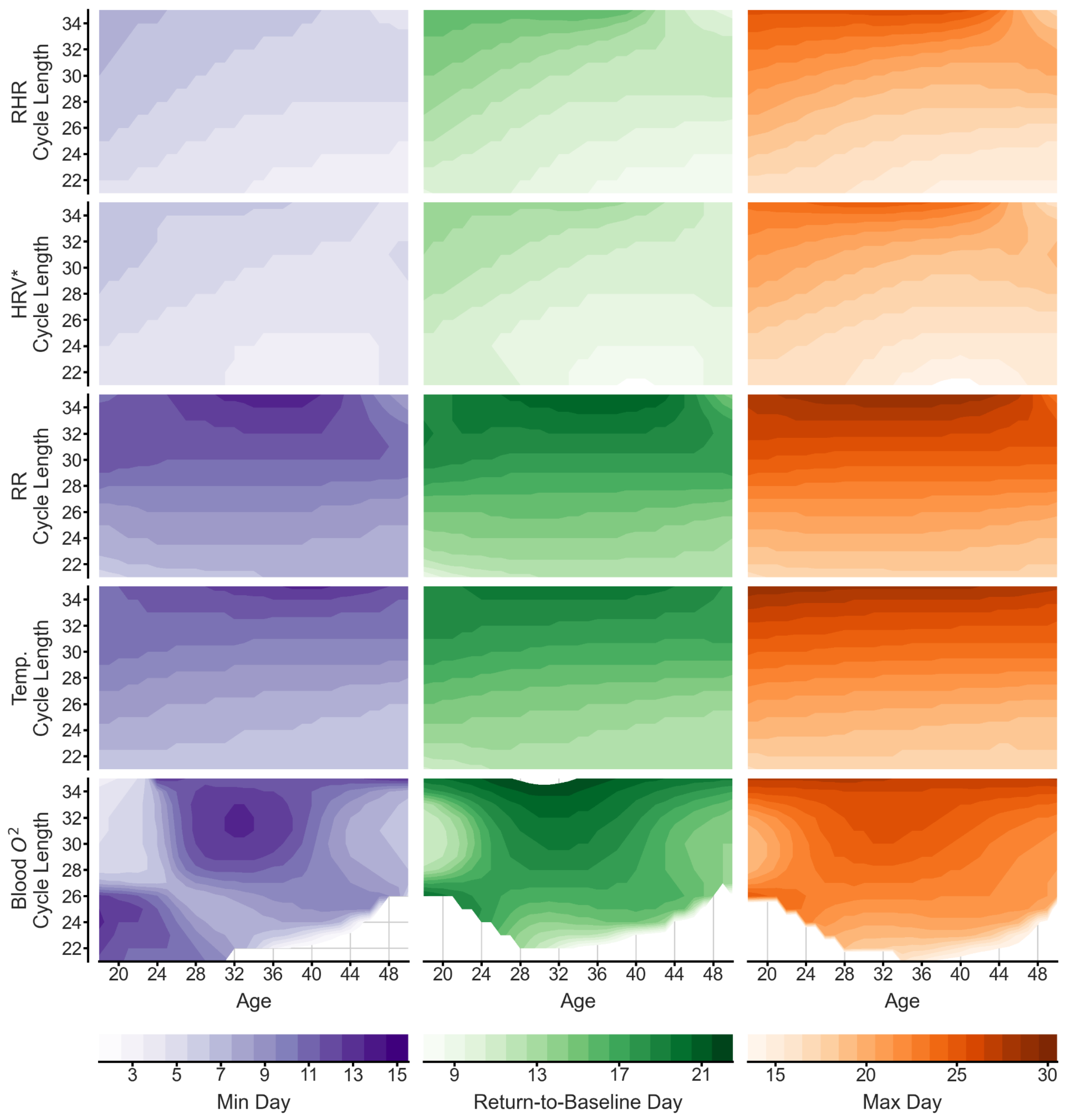


**Figure S8. Model-generated biometric critical values across age and cycle length.** We used Generalized Additive Models to generate temporal profiles at different ages and CL. With these models, we evaluated across combinations of age (x-axis) and cycle length (y-axis) to identify the day where each biometric reaches its minima, return-to-baseline, and maxima. Heat maps represent the day each biometric reaches each of these points post cycle onset (day 0).

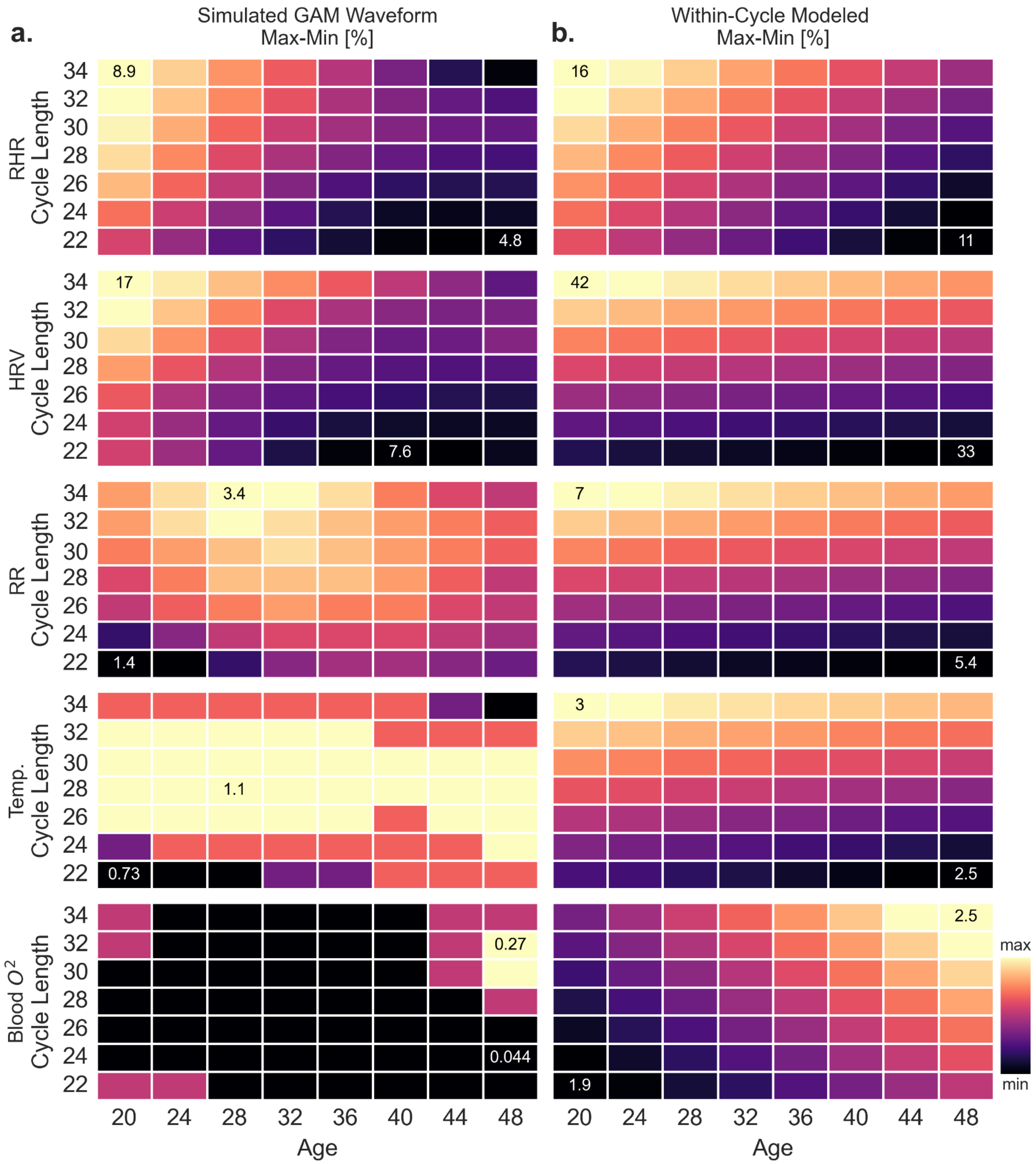


**Figure S9. Range of biometrics across the cycle for different ages and cycle lengths. (a)** The range (percentage difference between max and min) computed from the GAM-generated normative profiles. Each square on a heatmap represents the range for a given age and cycle length combination. Colors linearly scale by magnitude within a biometric heatmap. Annotations at light color indicate the maximal range across the combinations of age and cycle length, similarly the dark color annotation indicates the minima. The cyclicality of the GAM waveforms fixes the temporal relationship between a waveform's maxima and minima; thus, these charts reflect the range over the "typical" cycle. **(b)** The range (percentage difference between max and min) generated from a model fit to participant's within-cycle ranges. In this approach, the maximum and minimum of each biometric was extracted by cycle, then the range was modeled with a GEE to generate the ranges for the different combinations of age and cycle length. Without the temporal constraints of the GAM in (a), this approach can be more reflective of the expected biometric range for a menstrual cycle. Both approaches accounted for BMI, sleep, workout, and seasonal covariates.

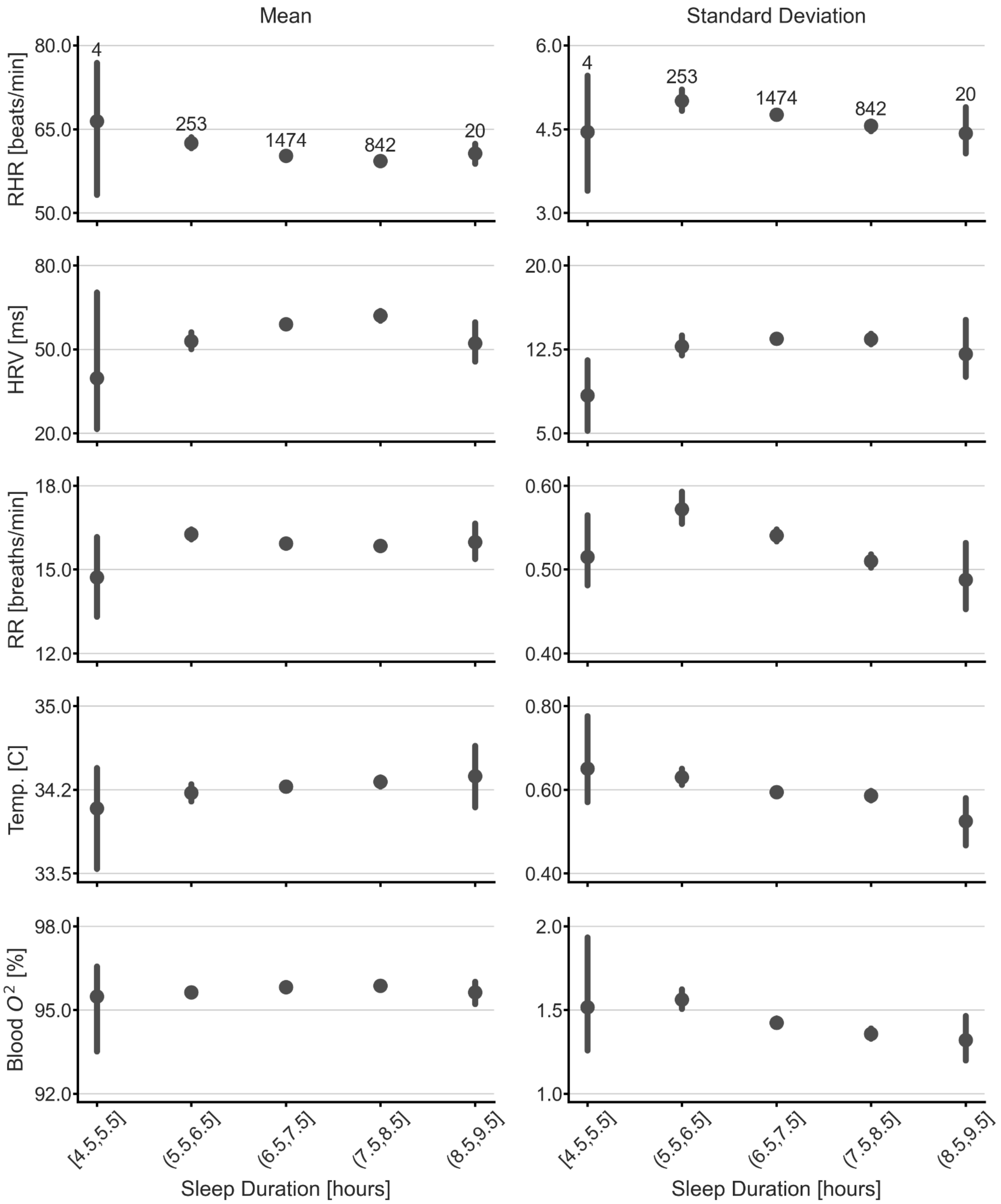


**Figure S10. Raw physiological biometrics by sleep duration.** Resting heart rate (RHR), heart rate variability (HRV), respiratory rate (RR), skin temperature (Temp.), and blood oxygen saturation levels (Blood O2) across sleep durations. Each point represents the (left column) mean value or (right column) standard deviation of that biometric across participants for a given 1-hour sleep duration bin from 4.5 - 9.5 hours (x-axis). Annotations above the points indicate the number of participants per bin. We did not examine trends in the extreme bins where we had limited participant data (i.e., equal to or fewer than 5.5 hours of sleep and equal to or greater than 8.5 hours of sleep).

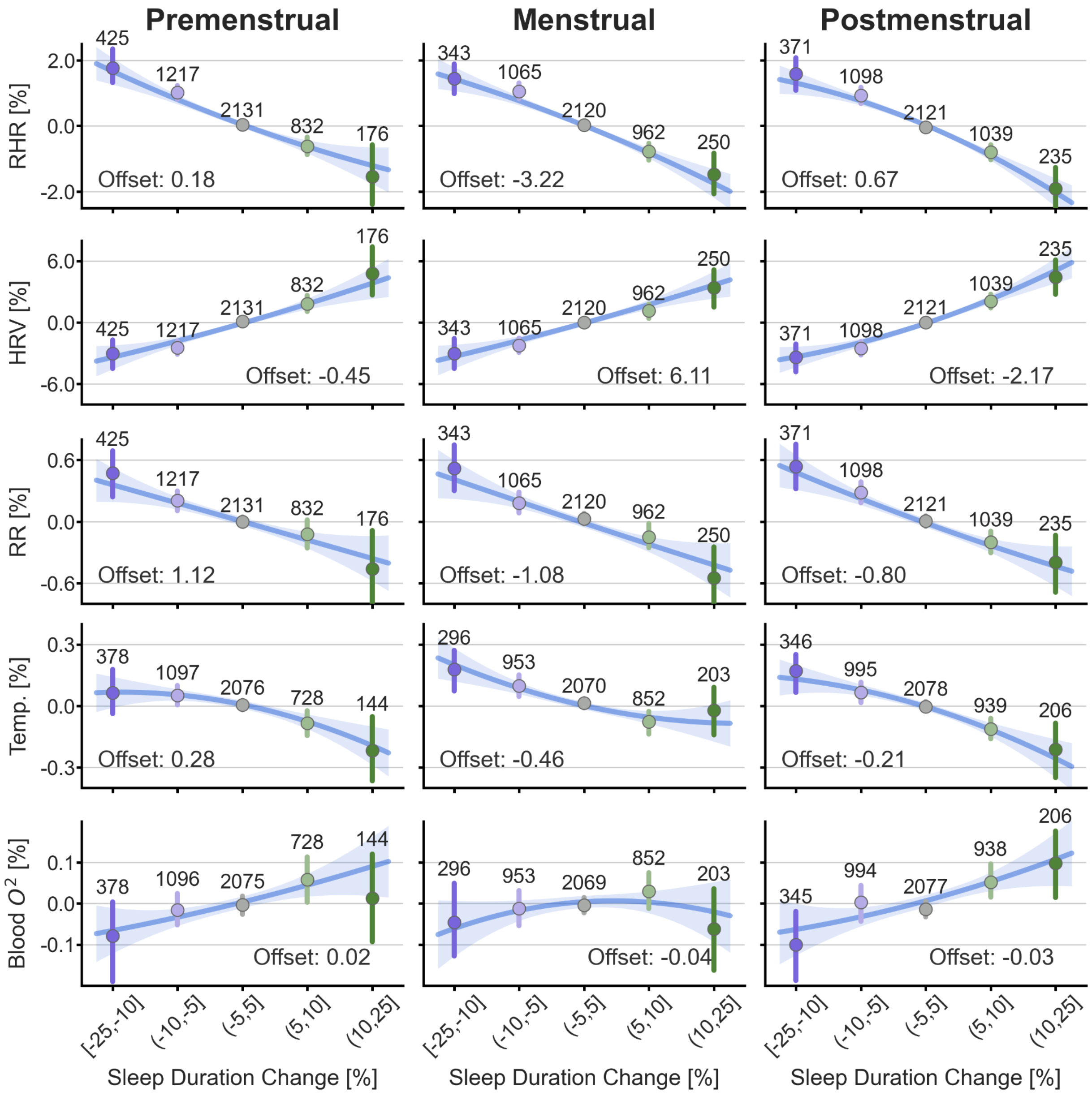


**Figure S11. Biometric responses to acute changes in sleep duration.** All biometric responses linearly scaled with changes in sleep duration across the menstrual cycle except for skin temperature. Points correspond to the average biometric on the last day of the cycle week when the sleep change occurred (day -1 for premenstrual, day 6 for menstrual, day 13 for postmenstrual). Annotations above each point represent the number of participants with at least one cycle at that sleep duration change, and error bars indicate the bootstrapped 95%CI of the mean. The blue line represents the predicted biometric at different sleep duration change levels, from a GEE model fit to the data, using the population mean of the covariates (age, BMI, sleep duration). We shifted the y-axis for each menstrual cycle week to reflect the change from the no-change in sleep duration condition (|change|<5%; the y-axis offset is annotated for each biometric).
